# Fever-like temperature impacts on *Staphylococcus aureus* and *Pseudomonas aeruginosa* interaction, physiology, and virulence both *in vitro* and *in vivo*

**DOI:** 10.1101/2023.03.21.529514

**Authors:** EC Solar Venero, MB Galeano, A Luqman, MM Ricardi, F Serral, D Fernandez Do Porto, SA Robaldi, BAZ Ashari, TH Munif, DE Egoburo, S Nemirovsky, J Escalante, B Nishimura, MS Ramirez, F Götz, PM Tribelli

## Abstract

**Background:** *Staphylococcus aureus* and *Pseudomonas aeruginosa* cause a wide variety of bacterial infections and coinfections, showing a complex interaction that involves the production of different metabolites and metabolic changes. Temperature is a key factor for bacterial survival and virulence and within the host, bacteria could be exposed to an increment in temperature during fever development. We analyzed the previously unexplored effect of fever-like temperatures (39°C) on *S. aureus* USA300 and *P. aeruginosa* PAO1 microaerobic mono- and co-cultures compared with 37°C, by using RNAseq and physiological assays including *in-vivo* experiments.

**Results:** In general terms both temperature and co-culturing had a strong impact on both PA and SA with the exception of the temperature response of monocultured PA. We studied metabolic and virulence changes on both species. Altered metabolic features at 39°C included arginine biosynthesis and the periplasmic glucose oxidation in *S. aureus* and *P. aeruginosa* monocultures respectively. When PA co-cultures were exposed at 39°C they upregulated ethanol oxidation related genes along with an increment in organic acid accumulation. Regarding virulence factors, monocultured SA showed an increase in the mRNA expression of the *agr* operon and *hld, pmsα* and *pmsβ* genes at 39°C. Supported by mRNA data, we performed physiological experiments and detected and increment in hemolysis, staphylxantin production and a decrease in biofilm formation at 39°C. On the side of PA monocultures, we observed increase in extracellular lipase and protease and biofilm formation at 39°C along with a decrease in motility in correlation with changes observed at mRNA abundance. Additionally, we assessed host-pathogen interaction both *in-vitro* and *in-vivo*. *S. aureus* monocultured at 39°C showed a decrease in cellular invasion and an increase in IL-8 -but not in IL-6- production by A549 cell line. PA also decreased its cellular invasion when monocultured at 39°C and did not induce any change in IL-8 or IL-6 production. PA strongly increased cellular invasion when co-cultured at 37°C and 39°C. Finally, we observed increased lethality in mice intranasally inoculated with *S. aureus* monocultures pre-incubated at 39°C and even higher levels when inoculated with co-cultures. The bacterial burden for *P. aeruginosa* was higher in liver when the mice were infected with co-cultures previously incubated at 39°C comparing with 37°C.

**Conclusion:** Our results highlight a relevant change in the virulence of bacterial opportunistic pathogens exposed to fever-like temperatures in presence of competitors, opening new questions related to bacteria-bacteria and host-pathogen interactions and coevolution.

## Background

*Pseudomonas aeruginosa* and *Staphylococcus aureus* are two opportunistic bacterial species globally distributed (1,2), which cause a variety of infections ranging from mild to severe, including skin and respiratory infections (3,4). Additionally, these bacteria are a major cause of both morbidity and mortality in cystic fibrosis patients, causing acute and chronic infections that lead to severe respiratory damage and (or perhaps due to) a continuous inflammatory response.

The interaction between *S. aureus* and *P. aeruginosa* has been recently reviewed (5) and it is traditionally described as antagonistic due to the secretion of different anti-staphylococcal factors by *P. aeruginosa*. Among these factors, the extracellular protease LasA affects the cell wall of *S. aureus*, leading to cell lysis and rhamnolipids cause biofilm dispersion^6^. *S. aureus* is sensitive to respiratory inhibitors secreted by *P. aeruginosa* such a*s* the pigment pyocyanin, quinoline N-oxides, and cyanhydric acid inhibit *S. aureus* respiratory chain affecting (6) the cytochrome bd quinol oxidase, which oxidizes ubiquinol and reduces oxygen as a part of the electron transport chain(7).

Despite this disadvantage, in co-culture with *P. aeruginosa*, *S. aureus* changes its metabolism to a fermentative state leading to the formation of small colony variants (SCVs). In this state *S. aureus* is resistant to pyocyanin and cyanide and able to survive in co-culture with *P. aeruginosa*. *S. aureus* cells can also shift to L-forms to escape the cell wall lytic activity of LasA(8,9). However, during the last years, the coexistence of both bacteria has also been detected in cystic fibrosis patients and *in vivo* experiments, revealing even a cooperative interaction(10–12). Although the interaction between both bacteria has been less explored under low oxygen conditions (9–11), microerobiosis is relevant in cystic fibrosis and other pulmonary diseases due to the presence of mucus and oxygen gradients (13). In this context, Pallet *et al.* (14) reported that anoxia leads to a phenotype of *P. aeruginosa* that is less dominant over *S. aureus*.

Besides oxygen and nutrients, another key factor for bacterial survival and physiology is temperature (15,16). Particularly, pathogenic species display virulence factors depending on several factors, including the host temperature (17,18). Previous studies have analyzed the effects of temperature transitions from the environment to the host on *S. aureus* and *P. aeruginosa* physiology by using transcriptomic and physiological approaches (19,20). In bacteria, the main temperature sensing mechanisms are RNA thermometers, which regulate the expression of genes at the post-transcriptional level through structured RNAs (18). RNA thermometers regulate different physiological features like the virulence factor *tviA* in *Salmonella typhi* or the type three injectisome in *Yersinia pseudotuberculosiss* (18,21,22). Temperature-dependent traits, mainly associated with virulence, host colonization and survival, have been recently reviewed in *Pseudomonas* spp., including *P. aeruginosa* (23).

In mammals, fever or pyrexia is defined as the increase in body temperature above homeostatic values, and, in the context of infections, is considered a beneficial process for the host (24,25). For example, in three different *Salmonella enterica* serovars, fever-like temperatures impair infective characteristics, suggesting that fever is a signal for a persistent state (26). In *S. aureus* USA300 strain AH1263, Bastok *et al.* (20) analyzed three different temperatures, including extreme pyrexia, and found alterations in some relevant phenotypes related to virulence. However, these authors focused mainly on the differences between 34°C and 37°C under aerobic conditions.

The aim of the present study was to analyze the effect of fever-like temperatures (39°C) on the interaction between *S. aureus* and *P. aeruginosa* by using a sequential culture scheme to analyze the RNA expression profile *in vitro* as well as virulence traits both *in vitro* and in a mouse model. Our data support that fever-like temperatures increase virulence *in vitro* and *in vivo*, particularly when *S. aureus* and *P. aeruginosa* are co-cultured.

## Results

### *S. aureus* and *P. aeruginosa* survival in co-cultures under microaerobic conditions and fever-like temperatures

We first investigated bacterial survival in microaerobic monocultures and co-cultures by using KNO_3_ as an alternative electron acceptor to oxygen. The cultures were incubated at 37°C either for 4 h (herein called 37°C) or for 2 h, followed by incubation at 39°C for 2 h (herein called 39°C) (**Fig. 1**). The CFU ml^−1^ value for *S. aureus* USA300 (herein called SA) in monoculture at 39°C was similar to that of the control cultures at 37°C (Fig. 2A), indicating that bacterial viability was not compromised by temperature in these conditions. Similar results were observed for *P. aeruginosa* PAO1 (herein called PA) monocultures (Fig. 2B). The same results -no differences-were obtained when assessing the effect of temperature on the survival co-cultured cells for both SA and PA (Fig. 2 A,B). When comparing mono and co-cultures, our results showed that under the conditions evaluated, PA did not show any differences when co-cultured (Fig 2 B.), while SA survival depend on the presence of the competitor, being significantly lower when co-cultured with PA and with no additional effects coming from temperature. Additionally, competence, determined by PA’s growth inhibition of SA, was also unaffected by temperature according to plate competence assays (Fig. 2C).

**Figure 1:**
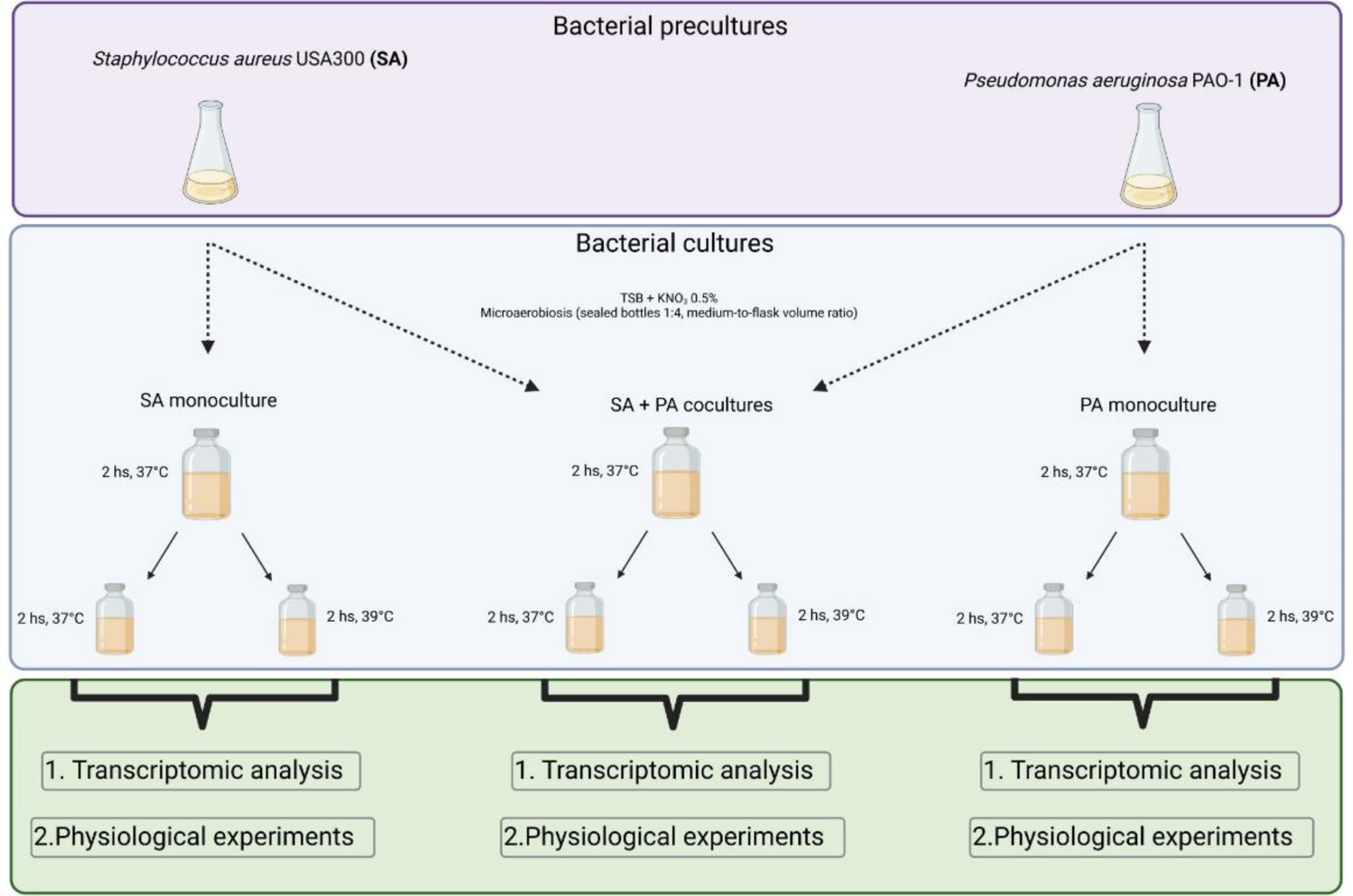
Culture conditions and experimental design. Experimental scheme to analyse the effect of fever-like temperatures on the physiology and interaction of *S. aureus* USA300 (SA) and *P. aeruginosa* PAO1 (PA).

**Figure 2:**
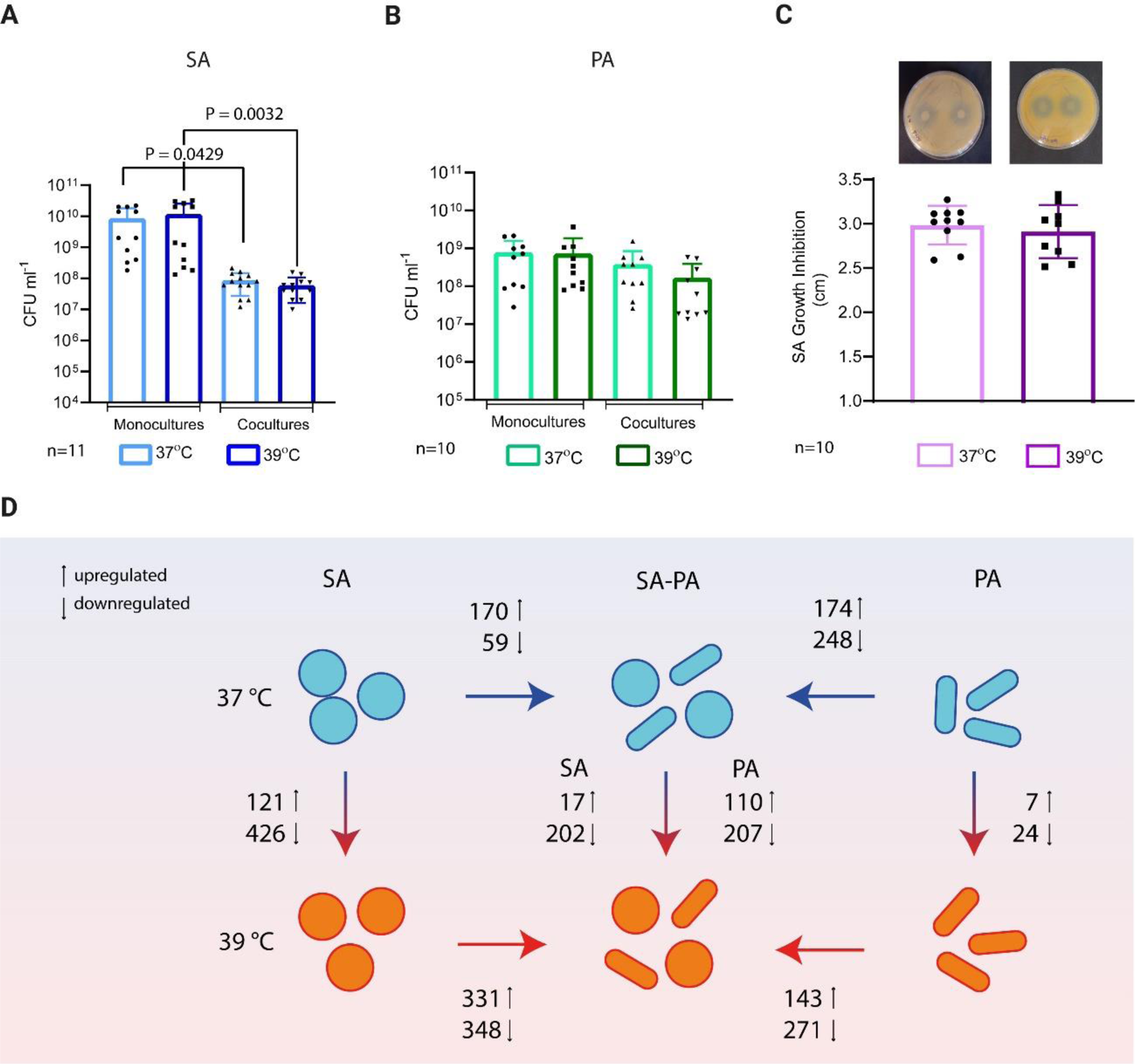
Survival and competence assays. SA and PA were cultured under microaerobic conditions either at 37°C or 39°C following the scheme shown in Fig. 1, in mono- or co-culture, and survival was measured as CFU/ml on selective media. **A.** SA count was determined on TSA+NaCl plates (1-way ANOVA, only statistically significant comparisons shown). **B.** PA was determined on cetrimide plates for PA (1-way ANOVA, non-significant). **C.** Competence test in agar plates performed as described in Materials and Methods (Unpaired t-Test, non-significant) **D.** Summary of the differential expressed gene number obtained by RNAseq experiments across different culture and temperature conditions.

### Gene expression profile of *S. aureus* and *P. aeruginosa* after exposure to fever-like temperatures in different culture conditions

In order to investigate how these bacterial species, adapt their physiology to fever-like temperatures, we next carried out total RNAseq experiments and physiological assays for SA and PA monocultures and SA-PA co-cultures, following the same culture scheme as that described above (Fig. 1). We analyzed the data using different approaches, including those included in Rockhopper software to detect differentially expressed genes (Fig S1 and S2), summarized in Fig. 2D and were further classified by cellular function (Fig S3 and S4 and Tables S1 and S2).

We performed pairwise comparison of the gene expression profiles for both PA and SA comparing the following conditions: SA or PA monocultures at 39°C and 37°C (herein called mono 39°C vs 37°C), SA or PA co-cultures and monocultured at 37°C (further called co vs mono 37), SA or PA co-cultures and monocultured at 39°C (further called co vs mono 39) and co-cultures at different temperatures (herein called co 39 vs 37). Overall, our results summarized in Fig. 2D showed a strong transcriptional change for most of the conditions. The only exception was monocultured PA that showed minimal change upon temperature treatments (only the 0.5 % of total expressed genes showed differential expression) while SA monocultures showed a strong response since we identify genes differentially expressed representing the 38% of the total expressed genes. However, when co-cultured, PA also shows a sharp response to fever-like temperatures when compared to monocultures (Fig. 2D).

Additionally, we performed a principal component analysis (PCA) (Fig. S5A and B). For SA, the PCA showed that the first PC discriminated culture condition (mono or co), while the grouping by temperature was less evident in any of the first three components (Fig. S5A), and the first and second components explained 89.8 % of the data variation (Fig. S5A). For PA, the first and second components explained 88.6 % of the data variation (Fig. S5B).

### Metabolic reshaping of *S. aureus* monocultures after exposure to fever-like temperatures

We also performed a Gene Ontology Enrichment (GOE) analysis, which allows the detection of a global trend of the metabolic pathway or branch overrepresented within the differentially expressed genes, even if not all the genes involved present significantly differences in the RNAseq analysis (Table S3 and S4). The GOE analysis in the SA mono 39°C vs 37°C expression dataset showed an enrichment in the arginine, tricarboxylic acid (TCA) cycle and fermentative pathways (Tables S1 and S3).

Particularly, in tryptic soy broth (TSB) cultures, we found that some genes related to arginine metabolism were upregulated at 39°C (Fig. 3A, Table S1, Table S5). To test if temperature affects arginine metabolism, SA monocultures were grown in complete defined medium (CDM) supplemented with glucose and KNO_3_ but without arginine or proline, following the same temperature scheme as that described above. Results showed that bacterial count in glucose-supplemented medium without arginine or proline was higher at 39°C than at 37°C (Fig. 3B), supporting the RNAseq observations and suggesting that arginine biosynthesis is triggered by fever-like temperatures.

**Figure 3:**
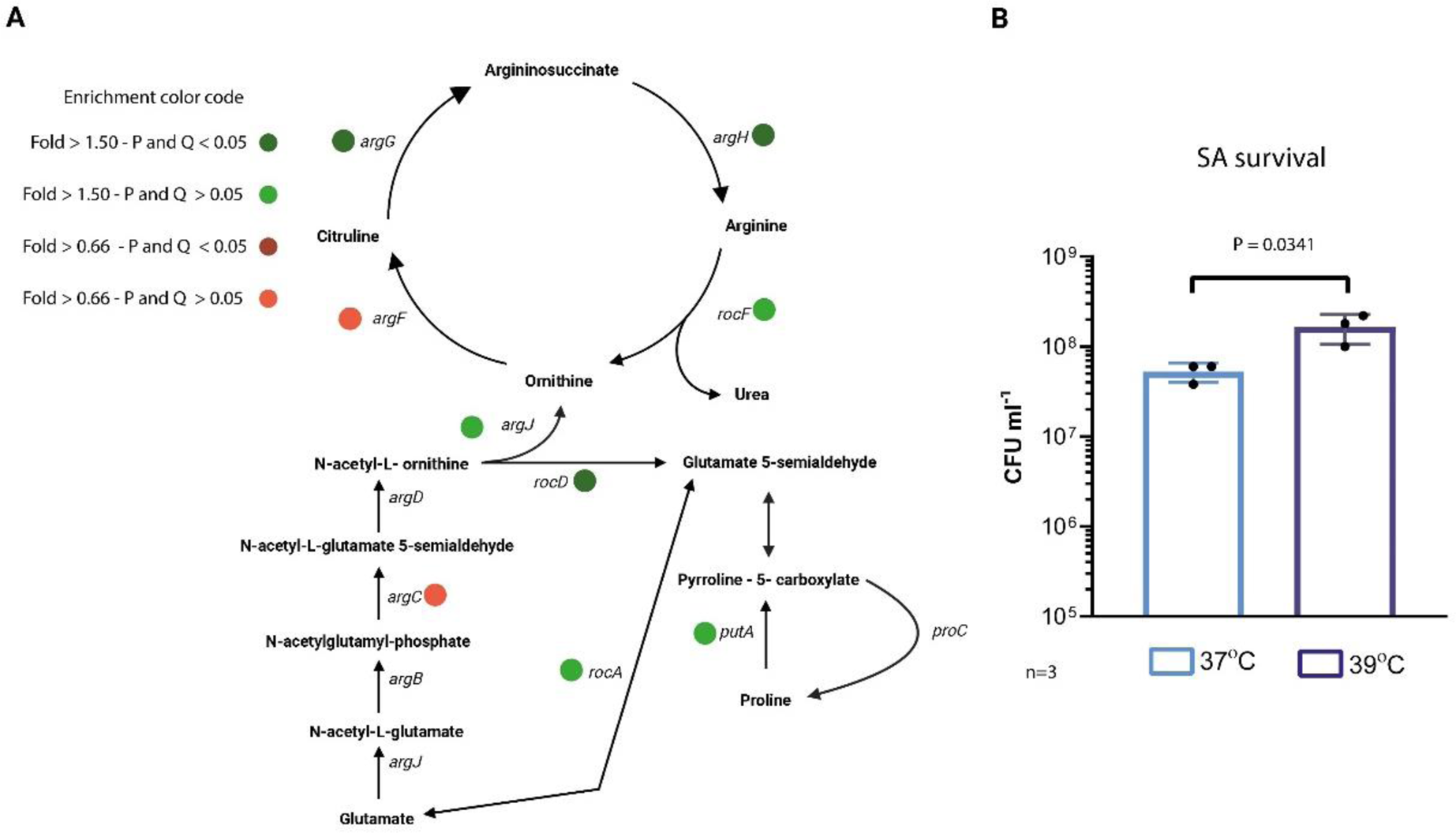
A. Effects of temperature and culture conditions in SA Arginine metabolism. Schematic overview of arginine biosynthesis and degradation metabolism in SA. Genes with changes at 39°C (both with a significant difference or a non-significant trend are indicated with color coded spots on the figure). **B.** SA survival in CDM medium lacking arginine and proline and supplemented with 0.5% KNO_3_. Cultures were incubated under microaerobiosis for 2 h at 37°C and further split and incubated at 37°C or 39°C for 2 h. CFU/ml was determined on TSA agar plates and compared by Unpaired t-Test.

Regarding the TCA cycle pathway, we found that genes were upregulated at 39° along with genes related with peripheral feeding pathways (Fig. 4A, Table S1, Table S5). On contrary, genes encoding proteins related to fermentation branches, were in general downregulated at 39° (Fig 4A, Table S1, Table S5) with the exception of the *fdhD* gene encoding the formate dehydrogenase subunit that was upregulated at 39°C. Also, the expression of the *fdh* and *fdhA* genes was increased, albeit non-significantly (Table S1, Table S5).

**Figure 4:**
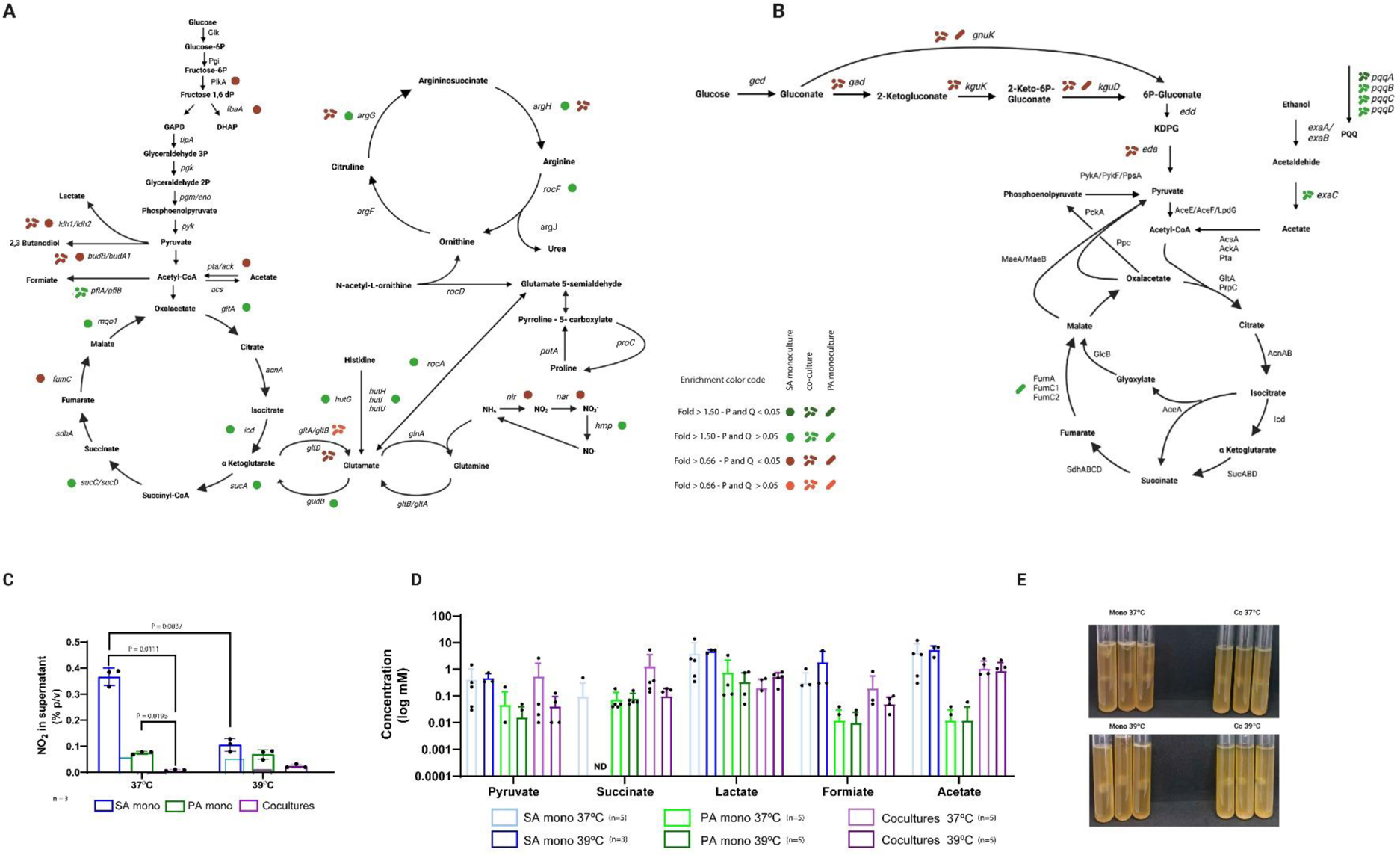
Effects of temperature and culture conditions in SA and PA central metabolic pathway. A. Schematic overview of SA carbon central metabolic pathways for monocultures 39°C vs 37°C and co-cultures 39°C vs 37°C. **B.** Schematic overview of PA carbon central metabolic pathways for monocultures 39°C vs 37°C and co-cultures 39°C vs 37°C.**C.** Nitrite production in SA and PA monocultures and co-cultures at 37°C and 39°C as described in Fig.1. Nitrite concentration in the supernatant was determined using the colorimetric Griess assay. Only significative differences are displayed with their corresponding P value after using 2-way Anova, Tukey’s multiple comparisons test. **D.** Organic acid content in the culture supernatant was determined by HPLC in CDM cultures supplemented with glucose and 0.5% of KNO_3._ Cultures were incubated following the scheme shown in Fig.1. **E.** Denitrification assay using Durham tubes. Monocultures and co-cultures were performed in TSB+ KNO_3_ (0.5%) and incubated at 37°C or 39°C for 48 hours and bubble size was observed.

Although the cultures were incubated under microaerobic conditions in the presence of nitrate, some genes encoding nitrate and the responsive nitrogen regulation system NreABC, which activates nitrate and nitrite dissimilation (27), were downregulated in mono 39°C vs 37°C along with nitrite reductase genes which catalyzed a reaction not coupled with energy production^27^ (Fig. 4A, Table S1, Table S5). In concordance with the gene expression profile, nitrite accumulation in the supernatant of SA monocultures was significantly decreased at 39°C as compared to 37°C (Fig. 4C) (27). In SA monocultures at 39°C the expression of cytochrome D ubiquinol oxidase coding genes was repressed, while the rest of the genes encoding cytochrome subunits were similarly expressed at 39°C and 37°C (Fig. S6A, Table S1, Table S5). Additionally, when we investigated organic acid production in SA monocultures in CDM supplemented with glucose at 37°C and 39°C under microaerobic conditions by high performance liquid chromatography (HPLC), we detected a decrease in succinate accumulation in the culture supernatant, but an increase in formiate at 39°C vs 37°C, while the rest of the organic acids were similar between temperatures (Fig. 4D). The integration of results suggests that SA monocultures feed the TCA cycle at 39⁰C through α-ketoglutarate to obtain energy, probably through cytochrome quinol and the succinate oxidase complex (Fig. 4A).

### *P. aeruginosa* gene expression profile in monocultures and metabolic features after fever-like temperature exposure

The few changes observed in PA monocultures after treatment with fever-like temperatures included an operon reported as iron responsive (28) that included *fumC1,* which encodes a fumarate reductase and *sodM* was upregulated at 39°C. On contrary, another key *Pseudomonas* metabolic pathway, the periplasmic glucose oxidation, was repressed at 39°C, with almost all the genes repressed (*kgut*, *kguk*, *kgD*, and *kgnt*) (Fig 4B, Table S2, Table S4, Table S6).

Nitrite production was similar for PA monocultures incubated at both temperatures (Fig. 4C) while in the Durham tube assay, gas production after 48 h was lower at 39°C, although growth was similar to that observed at 37°C (Fig. 4E). The organic acid profile in CDM supplemented with KNO_3_ and glucose showed a low concentration of organic acids in the supernatants of PA monocultures, particularly acetate (Fig. 4D). Cytochrome expression showed no significant differences between temperatures (Table S2, Fig. S6B).

### Culture condition impacts on bacterial physiology depending on temperature

For SA in co vs mono at 37°C, the GOE analysis showed five over-represented categories, including purine metabolism, amino acid biosynthetic and catabolic processes, siderophore biosynthesis, and heme transport (Table S3). The heme degradation and staphyloferrin B biosynthesis pathways were also represented among the upregulated genes in co-cultures, when compared with monocultures at 37°C (Tables S1, Table S3, Table S5). Particularly, enrichment in the biosynthetic amino acid pathways for L-isoleucine, L-valine and L-leucine were over-represented, with genes belonging this metabolism upregulated in SA co-cultures (Table S1, Table S3, Table S5). In microaerobic co-cultures SA displayed fermentative metabolism with genes encoding Ldh and Pfl coding genes upregulated in co-cultures (Table S1, Table S3, Table S5).

In the same comparison (co vs mono 37), PA genes involved in the peripheral fructose catabolic pathway were repressed (Table S2, Table S4, Table S6). Additionally, the GOE analysis showed enrichment in cysteine metabolism, with upregulated genes belonging to the assimilatory sulfonate reduction superpathway (, Table S2, Table S4, Table S6).

RNAseq analysis of co vs mono 39 for SA showed upregulation in several glycolysis-related genes (Table S1, Table S3, Table S5) and staphyloferrin biosynthesis genes. Interestingly, TCA-enzyme-coding genes and three succinate dehydrogenase subunits coding genes were repressed (, Table S1, Table S3, Table S5). Concomitantly, genes related with fermentation *pflA*, *pflB* and three genes encoding lactate dehydrogenase showed a strong expression increase ranging from 121 to 4-fold in co vs mono at 39 (Table S1, Table S5).

The PA expression profile in the same conditions (co vs mono 39) revealed repressed genes belong to the periplasmic glucose oxidation pathway (Table S2, Table S4, Table S6). In contrast, we found overexpression of genes belonging to glyxiolate shunt and of genes encoding the succinate dehydrogenase subunits (Table S2, Table S4, Table S6), suggesting that in PA, glyoxylate could feed the TCA cycle through succinate in co vs mono 39. Moreover, genes encoding a L-lactate dehydrogenase (that oxidizes L-lactate) and permease were upregulated in co vs mono independently of temperature, but with a sharper effect at 39°C (3.6 and 7-fold for mono vs co at 37 and 7 and 20-folds for co vs mono at 39 respectively) (Table S2, Table S6). In concordance with lactate consumption, we detected lower lactate levels in co-cultures supernatants comparing with monocultures (Fig. 4D).

### Fever-like temperatures affect *S. aureus* and *P. aeruginosa* gene expression profile in co-cultures

When we compare co-cultures at different temperatures (co 39 vs 37), for SA we observed at 39°C the repression of genes belonging to transcription and translation processes as well as envelope modification, and of the enterotoxin-coding genes *seq* and *sek* (Tables S1 and S3). In contrast, only genes comprising the PTS type II ascorbate-specific operon, were upregulated at 39°C (Table S1). In general, when PA was present, SA central metabolic genes were not affected by temperature. Some exceptions were *ldh*, *budB*, and the cytochrome D ubiquinol oxidase subunits, which were downregulated at 39⁰C (Fig. S6A, Table S1, Table S5).

For PA, co 39 vs 37 expression profile analysis showed changes in key metabolic features. Genes belonging to the ethanol oxidation pathway, which is a secondary metabolic branch present in *Pseudomonas* species for energy production through oxidation of ethanol (29), were upregulated at 39°C (Fig.4B, Table S6). These included genes belonging to the PQQ biosynthesis, an iron-containing aldehyde dehydrogenase, the cytochrome C and the response regulator *erbR* (Fig. 4B, Fig. S6B, Table S2, Table S6). Genes related to anaerobic metabolism were repressed in co-cultures at 39°C in comparison with 37°C, even when both were incubated under low oxygen conditions in the presence of nitrate (Table S2, Table S6). Accordingly, nitrite accumulation in co-cultures was similar at both temperatures and lower than in monocultures after 4 h of incubation (Fig. 4C). Gas accumulation in Durham tubes after 48 h showed a similar bubble size and similar growth when comparing co 39 vs 37, but smaller when compared with monocultures (Fig. 4E). Comparing co 39 vs 37 organic acid content in the whole supernatant pyruvate, succinate and formiate content was decreased (Fig. 4D).

### Temperature has a different impact on *S. aureus* but not *P. aeruginosa* depending on the culture conditions

We additionally performed a multifactorial ANOVA for each gene present in the RNAseq with false discovery rate (FDR) correction method to understand if there were interaction between the analyzed factors being temperature and culture conditions (mono or co). We found no statistical interaction between the culture condition and temperature for PA (Fig. S5D). In contrast, for SA, this analysis showed an interaction between these factors, importantly *agr* operon and three of its target genes, crucial for SA virulence, presented interaction between temperature and culture conditions (Fig. S5C; Table S7).

SA monoculture, the *agrABCD* operon was overexpressed at 39°C in comparison with 37°C (four-fold on average, Table S1). As expected, *agr* overexpression caused an increase in the expression of its target genes, including *psmβ1*, *psmβ2* and *hld* (Fig. 5A, Table S1). Additionally, other genes described to be under direct or indirect control of *agr* were found to be repressed (Fig. 5A, Table S1). When we compared the SA expression profile of co 39 vs 37, co vs mono 39 or co vs mono 37, we did not find differences in *agr* genes or its direct targets although showed different expression trend (Fig. 5A).

**Figure 5:**
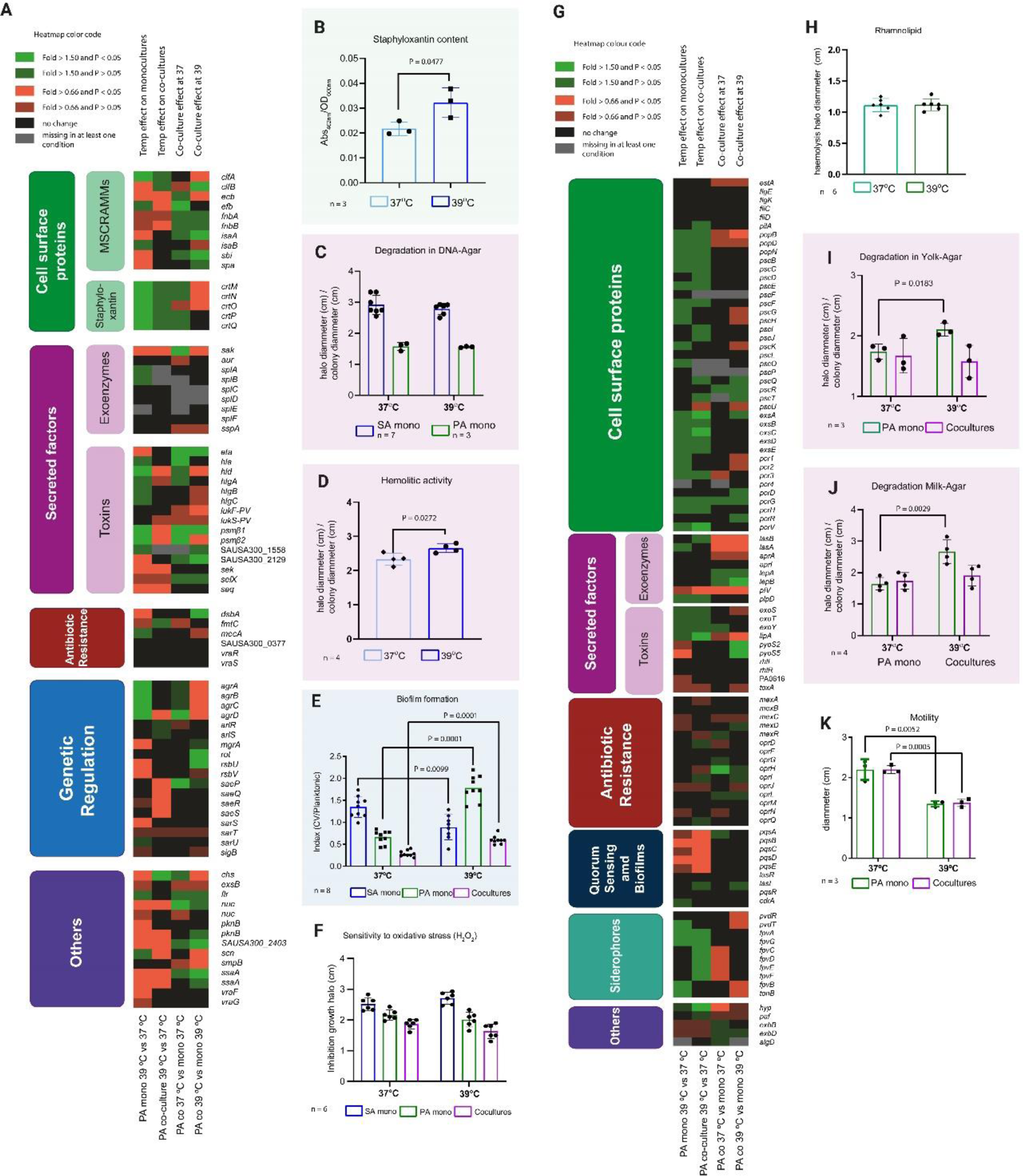
Fever-like temperatures trigger virulence factors and alter antibiotic resistance in both SA and PA. **A**. Heat map representing the gene expression profile of SA cultured in different conditions and temperatures. **B.** Staphyloxanthin production after incubating SA in monoculture following the scheme described in Fig. 1. Methanol-extracted pigment from supernatants was measured at 462nm **C.** Extracellular DNAse activity (degradation halo) for SA or PA monocultures assayed in DNA-agar plates incubated overnight at 37° C or 39° C. **D.** SA extracellular hemolytic activity tested in sheep blood-agar plates inoculated with SA monocultures incubated overnight at 37° C or 39°. Clear halo and the colony diameter are informed. **E.** Biofilm formation in multiwell plates with SA monoculture incubated overnight at 37° C or 39° C. Biofilm formation index was calculated as crystal violet (CV) ʎ_570 nm_/planktonic cells ʎ_595 nm_ for each replicate. **F.** Sensitivity to oxidative stress. Lawns of SA, PA or co-cultures were seeded in TSA plates. A 5-µl drop of H_2_O_2_ was placed in a sterile paper disk and further incubated at 37° C or 39° C. **G** Heat map representing the gene expression profile of PA cultured under different conditions **H.** Rhamnolipid production in PA monocultures was determined by the hemolytic halo in sheep-blood agar plates incubated at 37° C or 39°C. **I.** Lipase extracellular activity in egg-yolk-supplemented TSA plates incubated at 37°C or 39°C. Clear zones or white precipitates were determined. **J.** Protease extracellular activity in milk-supplemented TSA plates incubated at 37° C or 39°C. The clear zone and colony diameter were determined. **K.** Motility of PA mono- and co-cultures was analyzed in motility plates incubated at 37° C or 39° C. All diameter measurements were performed with the ImageJ software. Individual values for each measurement are shown. Unpaired t-Test was performed and only significant differences are informed with their corresponding P-value.

The SA transcriptional data related to virulence are summarized in Figure 5A. These data show that the expression profile of the master regulator system *agr* was affected by the fever-like temperature depending on the presence of the competitor and that the most virulent expression profile was found in SA monocultured at 39°C.

### Fever-like temperatures trigger virulence factors and alter antibiotic resistance in *S. aureus* and *P. aeruginosa in vitro*

Considering the impact of fever-like temperature (39°C) on gene expression, we tested its impact on some virulence factors of SA monocultures *in vitro,* including DNAse activity, pigment production, haemolysis, biofilm formation and colony phenotype in Red-Congo agar (30,31).

*In vitro* staphyloxanthin production was higher at 39°C, similar to what observed in the competence plates shown in Fig. 2C, in which the SA lawns presented a strong yellow color (Fig. 5B). Accordingly, the biosynthetic operon staphyloxanthin were overexpressed at 39°C (Figure 5A, Table S1, Table S5). Extracellular DNAse activity in DNA agar plates was similar at both temperatures after 24 h of incubation (Fig. 5C). We also tested the colony phenotype in Congo red plates, where SA presented similar, black-pigmented colonies both at 37°C and 39°C after 24 h of incubation. Haemolysis as tested in blood agar plates was higher at 39°C (Fig. 5D). Biofilm formation, as assayed in polystyrene multiwell plates using TSB medium supplemented with KNO_3_, was lower at 39°C, in line with our RNAseq data (Fig. 5E) and with the upregulation of *agr* under the same conditions (32).

Antibiotic resistance was screened using the Sensititre plate multi-test for *S. aureus* monocultures incubated at 37°C or 39°C during the entire test. We found that in general the antibiotic resistance was similar in SA monocultures, excepted for moxifloxacin and daptomycin that presented a 1-fold increase in the Minimal Inhibitory Concentration (MIC) at 39°C compared with those obtained at 37°C (Table S8). Additionally, we found no differences in the oxidative stress resistance assay for SA monocultures incubated at different temperatures. (Fig. 5F).

For PA mono 39 vs 37, RNAseq analysis related to the virulence factors summarized in Fig. 5G, showed that genes related to the quorum sensing *pqs* system were repressed at 39°C. In PA monocultures, extracellular DNAse activity, lipase and rhamnolipid production were similar at both temperatures (Fig. 5B, I and H). Extracellular protease activity of PA monocultures at 39°C was significantly higher while motility was lower (Fig. 5J and K). Antibiotic resistance at different temperatures was also tested using the Sensititre panel for Gram negative bacteria. We showed a decrease 2-fold in cefotaxime MIC and an increase in resistance to minocycline at 39⁰C (Table S9). Oxidative stress resistance showed similar results at both temperatures (Fig. 5F).

In co-cultures, the PA type III secretion system (TSS3) regulators *exsA* and *exsC* were upregulated at 39 as well as *pcrV* also a TSS3-related (Fig. 5G, Table S2). Other genes related with this system showed the same trend albeit non-statistically significant (Fig. 5G). The phenazine production genes and the extracellular lipase coding gene *lipA* followed the same trend, being upregulated at 39°C (Fig. 5G, Table S2). Considering these changes, we tested the extracellular lipase activity in co-cultures and observed higher activity at 39°C while extracellular protease activity showed similar results at both temperatures (Fig. 5I and J). Motility, which is mostly driven by PA, decreased at 39°C in co-cultures, similar to what was found in monocultures, thus showing a temperature effect despite the presence of SA. When we compared the expression of SA virulence factors at co 39 vs 37, we found repression at 39°C of the Sae operon *nuc, sek, seq, ecb, efb, pknB, scn, sak, agrD* and two direct targets of the *agr* system *psmβ1, psmβ2* (Fig 5A).

### Temperatures affects SA and PA induced cytokine production and invasion in human cell lines

To further investigate these *in vitro* observations, we analyzed cytokine production and cellular invasion on the epithelial lung cell line A549. Cells were infected with inoculum grown following the same culture scheme described in Fig.1. When A549 cells were infected with PA monocultures, production of the IL-8 and IL-6 cytokines was similar despite temperature (Fig. 6A and B). In contrast, when A549 cells were infected with SA monocultures at 39°C, proinflammatory IL-8 production was higher than that observed when cells were infected with SA monocultures at 37°C while IL-6 production was similar at both 37°C and 39°C (Fig. 6A and B). When cells were infected with SA-PA co-cultures, a similar production of IL-6 and IL-8 was observed regardless of the temperature, although a slight increment was observed for IL-8 production at 39°C (Fig. 6 A and B). Additionally, IL-6 production as lower when A549 cells were infected co-cultures comparing with PA monocultures at both temperatures (Fig. 6B).

**Figure 6:**
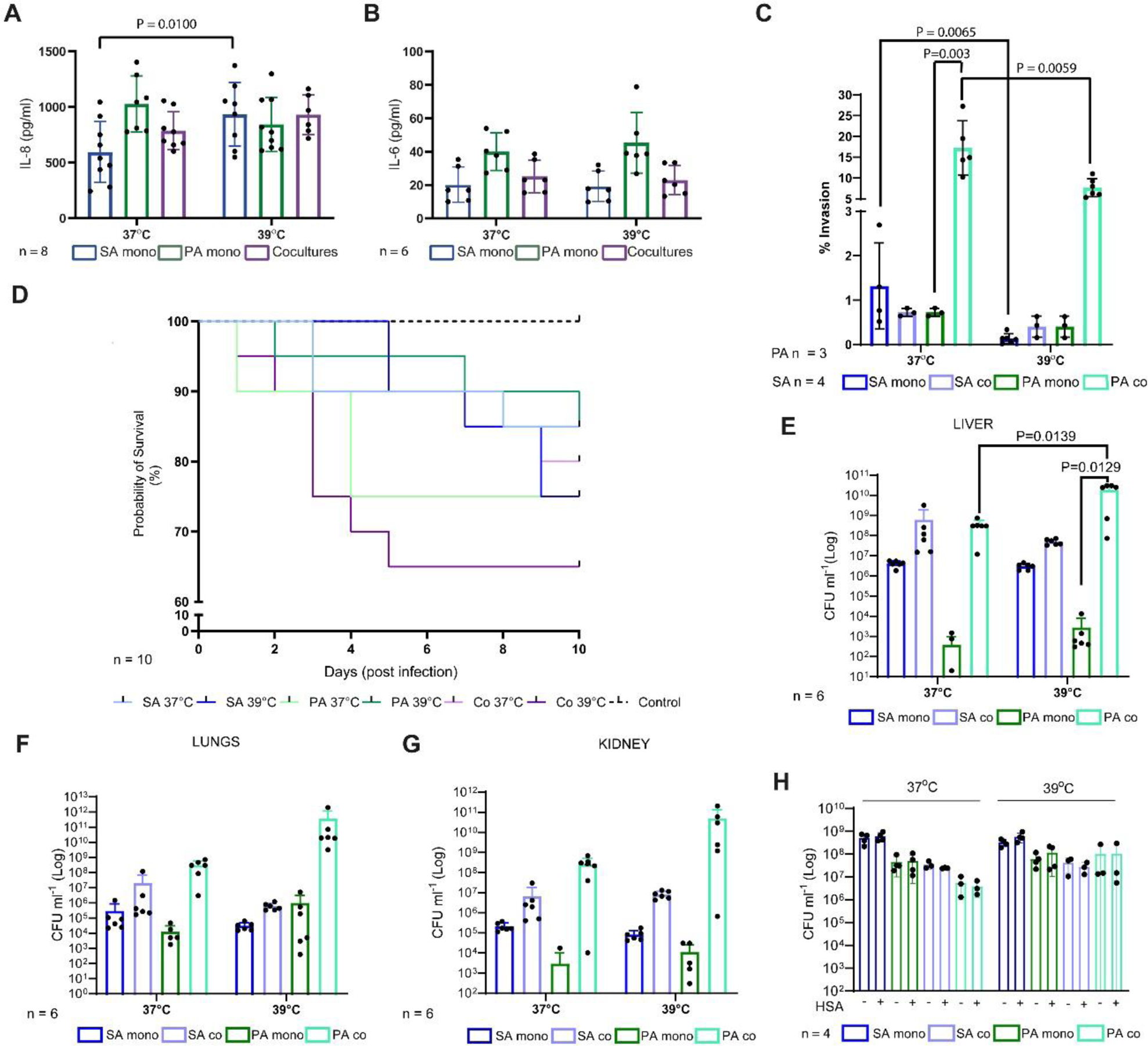
Fever-like temperatures impact on inflammatory cytokine production, cellular invasion and *in vivo* virulence. **A.** Production of IL-8 in the A549 cell line. **B**. Production of IL-6 in the A549 cell line. **C.** Invasion assay in A549 cell lines infected with bacterial cultures prepared as was shown in Fig.1. Results are presented as percentage of bacterial inoculum that was internalized. The results were analyzed for each microorganism separately. The graph displays two different scales for better visualization. **D.** Mouse survival was evaluated using 10 individuals for each group intranasally inoculated with mono- or co-cultures incubated as described in Fig.1. Sterile PBS was used as a negative control. Mouse survival was monitored daily for 10 days. The graph has a scale break for better data visualization. **E.** Bacterial burden in the liver. **F.** Bacterial burden in the lungs. **G**. Bacterial burden in the kidney **H**. Effect of human serum albumin on SA and PA survival at 37°C or 39°C. Zero values obtained in the bacterial count are not shown on the Log scaled graphs but mentioned on the graph and were considered for statistical analysis. Zero values included (non-infected control n = 6 for all organs; PA mono 37 n=1 in liver; n=1 in lungs and n=5 in kidney; PA mono 39° C presented zero values in n=1 in kidneys). A-C, H 2-way ANOVA E-G 1-way ANOVA, Sidak’s multiple comparisons test. Only significant differences displayed with their corresponding P-value for all the comparisons.

When the cellular invasion assay was performed with SA monocultures incubated at 39⁰C, a decrease in cellular invasion was observed, in agreement with a more virulent profile (Fig. 6C). In contrast, cellular invasion for SA in co-cultures no differences were found between temperatures (Fig. 6C). However, for PA in co-cultures we found a decrease in cellular invasion at 39°C comparing with 37°C (Fig. 6C). Overall, our results show that fever-like temperature affects SA virulence, suggesting an acute infective state when PA is absent while for PA the effect is the contrary.

### *In vivo* virulence is affected by temperature and culturing in a mouse model

We next performed intranasal infections in mice with bacteria cultured as we described in Fig.1. We used this simplified model in which the bacteria were previously exposed to different temperatures to understand how this factor impacts on bacterial virulence. We tested both mouse survival and bacterial burden in the mouse lungs, kidneys, and liver. Although there were not significant differences at the end of the experiment, we observed that survival was lower when mice were infected with SA monocultures preincubated at 39°C vs 37°C (Fig. 6D). In contrast, when mice were infected with PA monocultures, mouse survival was lower when the inoculum was grown at 37°C (Fig. 6D). Moreover, SA-PA co-cultures incubated at 39°C caused greater mortality than those incubated at 37°C and was the treatment that caused the highest mortality in mice (Fig. 6D).

Bacterial burden in the lungs, kidneys, and liver showed that SA count in these organs was similar when comparing between temperatures but was slightly higher when SA was in co-culture (Fig. 5 E, F and G). For PA we showed that in liver bacterial burden was significantly higher when we compared co-cultures and monocultures at 39° (Fig. 6 E, F and G). Importantly, PA co-cultures led to significantly higher counts in liver at 39⁰C than at 37°C and in kidney and lungs we also observed the same trend although the differences were not significant (Fig. 6 E, F and G).

Serum albumin is a major protein in plasma that affects gene expression, virulence and survival in *Acinetobacter baumanii* (33,34). To understand whether human serum albumin (HSA) could alter the bacterial number in co-cultures at 39°C, we grew the bacteria following the same culture scheme (Fig.1) but in the presence or absence of this protein. HSA did not affect SA or PA, which was expected since survival was not affected by 39°. In addition, we found no difference in the CFU count of SA or PA co-cultures at 37°C or 39°C in the presence or absence of HSA. However, as these experiments were carried out *in vitro* we cannot rule out an effect of HSA *in vivo*.

Finally, we wondered whether longer co-culture times could cause a different physiological response at 39°C in comparison with that observed at 37°C. Thus, we cultured SA, PA and PA-SA at 37°C and 39°C in the same microaerobic conditions but for 24 h. Surprisingly, co-cultures showed an evident, blue/green-colored medium typical of pyocyanin production but only at 39°C (Fig. S7A). More intriguing is the similar CFU count in the co-cultures after 24 h at both 39°C and 37°C, despite the different pyocyanin content (Fig. S7B).

## Discussion

The infection process and outcome depend on several factors, including the availability of nutrients, the microenvironment, the organ structure and the immune host response (35–37). Temperature is a physical factor that affects cellular physiology in the environment and during infection(38). Pyrexia or fever is the increase in the body temperature above a specific set-point controlled by the hypothalamus (39). This increase is often due to infection or other non-infectious causes such as inflammation or malignancy (39). In humans, fever-like temperatures have been classified as low-grade (37.3°C to 38.0°C), moderate-grade (38.1°C to 39.0°C; used in this work), high-grade (39.1°C to 41°C), and hyperthermia or hyperpyrexia when the temperature increases above 41°C (40). Fever is considered a beneficial process to resolve infections and increase host survival, whereas antipyretic treatments have been shown to increase mortality in influenza patients (41,42). Evans et al. (2015) reviewed the ways in which fever contributes to control infections, including direct effects like the increase in Gram-negative bacterial lysis caused by serum components and stimulation of the adaptive and innate immune system(42). However, host-pathogen coevolution implies the selection of bacterial features that counteract fever effects.

The effect of temperature has been studied mostly regarding the shift from the environment to the host. In *P. aeruginosa* PAO1 as well as in PA14, for example, genes encoding virulence factors have been found to increase at 37⁰C as compared with 22°C, particularly genes related to quorum sensing, exoproteins, and siderophores (19,43). For SA, Bastock et al. (2021) reported an expression profile rearrangement in cultures incubated at 40°C using aerobic monocultures (20). However, the authors performed human nasal epithelial cell line infection experiments comparing only SA incubated at 34°C and 37°C. In our model, we chose a sequential 2-h incubation at 37°C followed by 2 h at 39°C to understand the early response of bacteria to the temperature shift from the host to a fever-like situation under microaerobic conditions. Recently, Hamamoto et al. (2022) performed an *in vivo* RNAseq of *S. aureus* colonizing the mouse liver and reported that genes related to low oxygen levels are upregulated at 24 and 48 h post-infection(44). This shows the importance of microaerobiosis during infection, as that used in this work.

In monocultures, our results showed a different response of *P. aeruginosa* PAO1 and *S. aureus* USA300 to temperature. SA monocultures showed a decrease in nitrate reduction, decrease in the expression of genes related to fermentative metabolisms and the overexpression of arginine biosynthesis and TCA-related genes. Reslane et al. (2022) reported that when glucose and arginine are absent, SA is a functional prototroph(45). These authors also showed that in the presence of glucose, and even in the absence of arginine, *S. aureus* behaves like an arginine auxotroph showing that absence of arginine alone is not enough to trigger arginine biosynthesis. The authors suggested that, during abscess infection, where active macrophages are found, arginine and glucose are depleted, thus activating arginine biosynthesis from proline (27). Our hypothesis is that incubation of SA at a fever-like temperature for 2 triggers some physiological responses similar to those displayed after immune system activation even though glucose is present.

The most important SA response to 39°C under low oxygen conditions was the increase in the expression of the *agr* operon and the differential expression of its direct and indirect targets. In concordance with the work by Bastock et al. (2021) (20) performed with aerobic cultures, our *in vitro* virulence assays for SA showed staphyloxanthin production, but, in contrast to these authors, we did observe a significant increase in genes related to its biosynthesis. Another important feature for virulence is hemolysis. In this work, we showed upregulation of hemolysin-coding genes at 39°C and an increase in hemolysis after overnight incubation at 39°C. Contradictory results regarding this issue can be found in the literature. Bastock et al. (20) showed that when plates are incubated at 40°C for 30 minutes, hemolysis increases with temperature, in line with our results, while Palela et al. (46) reported that at 40.5°C, hemolysis decreases because temperature affects hemolytic pore kinetics. Thus, results of this and other works show the complex scenario when virulence factors are evaluated.

SA co-cultured with PA showed a strong increase in genes encoding fermentative enzymes and a repression of TCA. Although a fermentative metabolism for SA was expected in the presence of PA, the effect at 39°C was even sharper than at 37°C.

Here we reported for SA the increased in the expression of genes belonging to the heme degradation and staphyloferrin biosynthesis pathway in co vs mono under low oxygen conditions. These pathways are related to iron acquisition from heme, the most abundant iron source in the host and a key feature in the competition with PA (47). The increase also was observed despite temperature with a shaper increase at 39°C.

In contrast, PA monocultures at 39°C comparing with 37°C, showed a different pattern with a more robust metabolism, with less differentially expressed genes in response to fever-like temperatures and some virulence features increased at 39°C in agar plates.

Remarkably, L-lactate dehydrogenase and L-lactate permease were upregulated in co vs mono despite the temperature with a more pronounced effect at 39°C according with an increased expression of *ldh* gene in SA in co-cultures that was also higher at 39C. Although the PA consumption of lactate produced by SA was reported our results showed that under low oxygen conditions this metabolic relationship was also observed and that at 39°C this pattern was more marked(48).

When we compare co-cultures at 39°C and 37°C we showed a different expression pattern, with few upregulated genes in SA, only genes comprising the PTS type II ascorbate-specific operon, which has been reported to alter the properties of SA cell wall and thereby modulate virulence (31) were upregulated at 39°C. Interestingly, PA in co-cultures display a large number of differentially expressed genes between temperatures, being the most remarkable metabolic feature was the upregulation of genes related to ethanol oxidation that could be related with the increased fermentative profile of SA in co-cultures which was more pronounced at 39°C than at 37°C. The ethanol oxidation pathway was described for PA even at low oxygen tension (49)

Besides the metabolic pattern, we found an effect of SA monocultures incubated at 39°C on IL-8 production, which was increased at 39°C along with a decrease in cellular invasion according to the upregulation of *agr* operon. Chekabab et al. described an inhibitory effect of SA filtrated supernatants on IL-8 production in the airway epithelial cell line Beas-2B stimulated with *P. aeruginosa*(50). In this work, we showed a slight increase in IL-8 production in co-cultures at 39°C in A549 epithelial lung cells along with an increase in cellular invasion independent of culture temperature. Alves et al. reported an intermediate interleukin production by cells infected with mixed biofilms of *S. aureus* and *P. aeruginosa* compared to mono-species biofilms(51). Additionally, in a wound model these authors described an impairment in the wound closure along with a proinflammatory response when mixed biofilms were used (51). These results, along with our *in vitro* experiments, with lower invasion in A549 cells and IL-8 suggest an increased virulence of SA when was exposed to 39°C.

Although mouse is physiologically different from human it is still a widely used model. In mice hypothermia can be display but also temperature rise upon infections, LPS administrated in low doses and endotoxin induced inflammation have been reported (52,53) (Oka et al., 2003; Shiraki et al, 2021). Eskilsson et al (2021) have recently reported that, in mice, the humoral response is implicated in eliciting fever responses during localized inflammation(54). Due to the complex process of fever in mouse, we used a simplified *in vivo* model in which only the bacteria were exposed to fever-like temperature to understand the early physiological response and its consequences. Our results revealed that bacteria pre-incubated at 39°C showed a different behavior depending on the species and the culture condition (summary in Fig. 7). In monocultures albeit not significant differences, a different trend was observed for SA and PA at 39°C comparing to 37°C°. SA monocultures intranasally inoculated in mice caused higher lethality at 39°C, with similar bacterial burden in the different organs. In contrast, PA monocultures caused slight lower mouse mortality.

**Figure 7:**
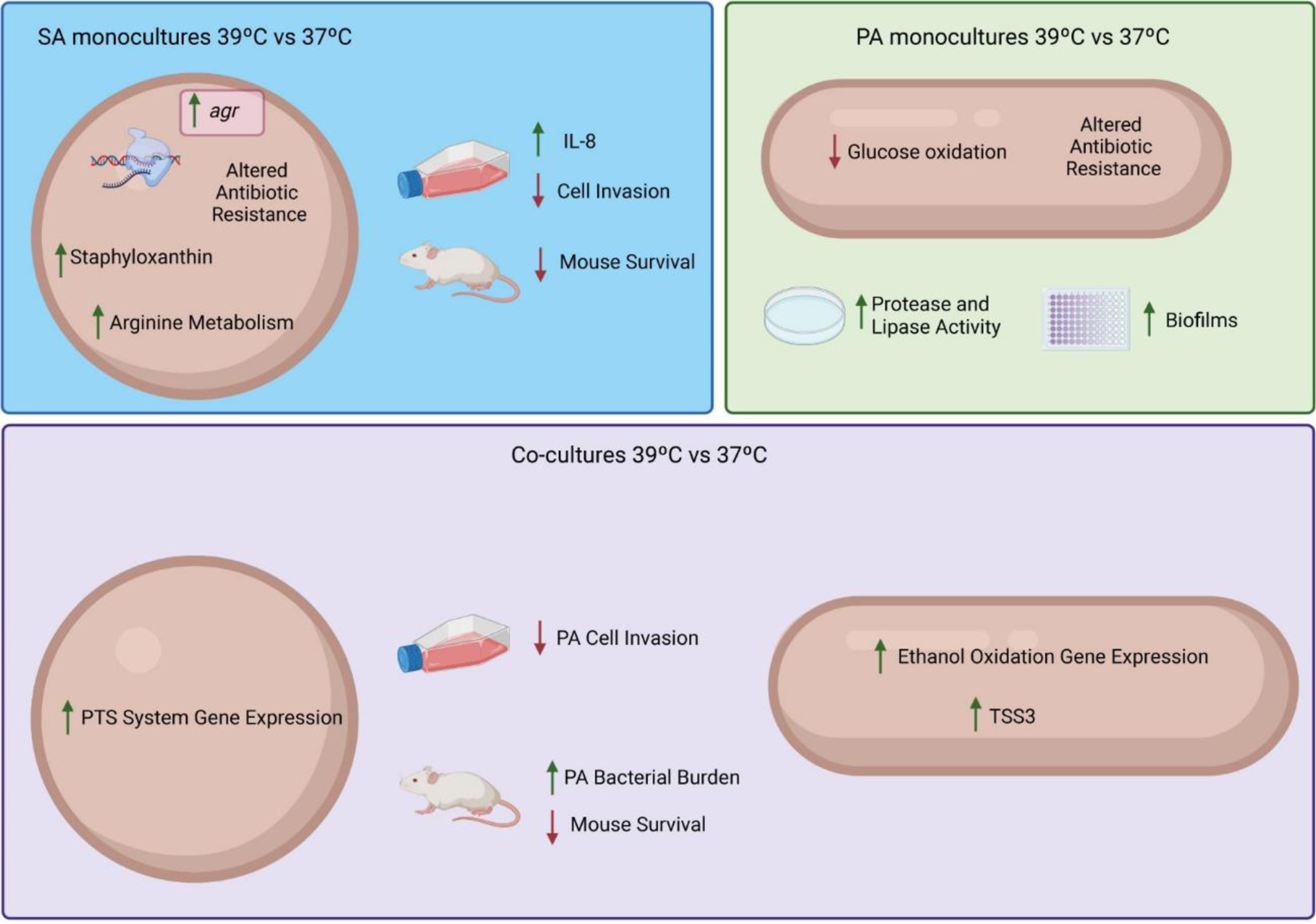
Summary of the relevant effects of temperature on *S. aureus* USA300 and *P. aeruginosa* PAO monocultures and co-cultures.

However, the most evident *in vivo* effect was observed for co-cultures incubated at 39°C. These severely impacted on mouse survival rate and led to an increase in PA burden in all the organs tested with significant differences in liver. In contrast, for SA the results were similar and independent of the temperature but higher than those obtained with monocultures, in agreement with the cell line invasion assays. In a keratinocyte cell line, Alves et al. demonstrated that *S. aureus* allows attachment of *P. aeruginosa* to cells, indicating a complex *in viv*o interaction between these bacterial species(51). In this work, we also observed that an overnight incubation of co-cultures at 39°C lead to a blueish color, typical of pyocyanin production but with a SA CFU count similar to 37°C co-cultures which did not show this color. Previous reports showed that pyocyanin production is lower under microaerobiosis comparing with aerobic conditions leading to a higher coexistence with SA (12). Our results are similar at 37°C to this previous report, however, at 39°C pigment production was observed even under microaerobic conditions. This increased production could increase virulence of PA at 39°C.

The mechanism by which these bacterial species sense the increase in 2°C and cause the phenotypes observed remains unknown. Thermoregulation through RNA thermometers has been described for SA but mostly for the environment-host transition and includes the *cspB*, *cspC*, and *cidA* genes (55,56). However, to our knowledge, the response to fever-like temperatures is still unknown. In PA, the sensing of temperature changes by RNA thermometers has also been reported for the virulence factors that are under the control of the quorum sensing regulator RhlR, including pyocyanin, elastase and rhamnolipid production but, again, the differences were observed following a temperature shift from 30°C to 37°C (57).

In summary, our results demonstrate that fever-like temperatures affect the virulence of *S. aureus (*SA) monocultures and *P. aeruginosa (*PA) and *S. aureus* co-cultures, mouse survival and bacterial burden in murine organs in co-cultures. Our results open new questions about the effect of fever-like temperatures on other pathogenic bacterial species and on the host-pathogen interaction.

## Methods

### Strains and culture conditions

The strains used throughout this work were *Staphylococcus aureus* USA300 (USA300_FPR3757) and *Pseudomonas aeruginosa* PAO1. Precultures were performed under aerobic conditions at 37°C in Tryptic Soy Broth (TSB; Oxoid). Microaerobic cultures were grown in TSB supplemented with 0.5% KNO_3_, in sealed bottles, using a 1:4 medium-to-flask volume ratio without agitation. SA and PA monocultures were inoculated at an OD_600nm_ of 0.05 and incubated at 37°C for 2 h. After that, cultures were split in two and further incubated at 37°C (control) or 39°C (fever-like temperature) for 2 h. A similar protocol was used for co-culture experiments, where SA and PA were co-inoculated each at an OD_600nm_ of 0.05.

### Plate competence assay

An overnight SA culture incubated at 37°C was adjusted to an OD_600nm_ value of 1 and then seeded on a Tryptic soy agar (TSA) plate. A 5-µl drop of a PA culture with an OD_600nm_ adjusted to 10 was placed in the center of the plate. After drying, the plates were incubated overnight at 37°C or 39°C. Statistical analysis: Unpaired t test.

### RNA extraction and RNA library preparation

SA and PA monocultures or co-cultures were incubated at either 37°C or 39°C, following the experimental scheme shown in Fig. S1. Frozen bacterial pellets were homogenized with milliQ water by using mortar and pestle devices in Eppendorf tubes. After cell disruption, total RNA was extracted with the Trizol method followed by extraction using a commercial kit (Total RNA extraction kit - RBC bioscience). Samples were treated with DNAse I (Promega). To improve the quality of the readings, ribosomal RNA was depleted from the samples by Novogen services (CA, USA). Libraries were constructed by Novogen Services (CA, USA). Mass sequencing was performed using the Illumina Novaseq 6000 platform with a paired-end protocol (Novogen Services; CA, USA). For each condition, triplicate independent RNA extraction and libraries were used.

### RNA-seq data analysis

Reads were preprocessed using the Trimmomatic computer tool(58) by eliminating adapters and low-quality sequences. The reads quality was evaluated using the FastQC tool (www.bioinformatics.babraham.ac.uk/projects/fastqc/).

Reads alignment and assembly, transcript identification and abundance quantification were carried out using the Rockhopper software for both monocultures and co-cultures(59). The reference genomes were *P. aeruginosa* PAO1 (AE004091.2) and *S. aureus* (USA300_FPR3757). Reads were normalized per kilobase per million mapped reads. To verify concordance between the independent replicates for each condition, a Spearman correlation analysis of normalized counts was performed. Differential gene expression was considered only with *P* < 0.05 and *Q* < 0.05 and a Fold Change (FC) ≤ −1.5 and ≥ 1,5. q-values are False Discovery Rate (FDR) adjusted p-values, using the Benjamini-Hochberg method. In the case of co-cultures, the RNA dataset was analyzed, and the reads aligned by separately using the PA or SA reference genomes.

Genes were sorted into functional classes using the KEGG (60), MetaCyc (61) and String (62) tools. For quantification and classification, tRNA transcripts were not taken into account.

### Data availability

RNA-seq data is available in the European Molecular Biology Laboratory (https://www.ebi.ac.uk/) under accession number E-MTAB-12581 (https://www.ebi.ac.uk/biostudies/arrayexpress/studies/E-MTAB-12581?key=6e13e32b-cae0-4154-9e08-1ccfa6fd46c8).

### Cultures and determination of organic acid production

To determine organic acid production, bacteria were grown in CDM (63) supplemented with 7.5 mM glucose as a carbon source. Cultures were centrifuged at 13,000 rpm for 5 min and supernatants were diluted 1:5 in water and filtered through 0.22-μm syringe filters (MSI, USA). Samples were analyzed by high performance liquid chromatography (HPLC) (LC-20AT Prominence; Shimadzu Corp., Japan) equipped with a UV detector (SPD-20AV; Shimadzu Corp.) using an Aminex HTX-87H column (Bio-Rad Laboratories, USA) at 50°C. The mobile phase consisted of 5 mM H_2_SO_4_ with a flow rate of 0.6 ml/min. Detection was performed at 210 nm and analytical standards (Sigma-Aldrich Co., USA) were used for quantification by external calibration curves.

### SA growth in arginine and proline free medium

To assess arginine metabolism in SA, bacteria were grown in arginine- and proline-free modified CDM or in CDM supplemented with glucose 7.5 mM. SA was inoculated at an OD_600nm_ of 0.05 and cultures were incubated at 37°C or 39°C under microaerobic conditions, following the same scheme as described in Fig. S1. After 4 h of incubation, the CFU ml^−1^ in TSA was determined.

### *In vitro* determination of virulence factors

The carotenoid pigment staphyloxanthin was quantified as previously described (64). Briefly, cultures grown as described in Fig. S1, the OD_600nm_ was adjusted to 1, and then 10 ml was centrifuged at 8000 rpm for 10 min. Cell pellets were washed with sterile PBS and resuspended in 1 ml of methanol. The tubes were incubated overnight in the dark with agitation at 37°C. Then, the tubes were centrifuged to collect the supernatant containing the extracted pigments. Staphyloxanthin was quantified by measuring absorbance at 462 nm. For the evaluation of virulence factors, precultures were grown at 37°C under aerobic conditions and the OD_600nm_ was adjusted to 1 for monocultures and for co-cultures in a 1:1 proportion for each bacterium, reaching an OD_600nm_ value of 1. With this bacterial suspension, a 5-µl drop was placed in the corresponding plates and the plates were incubated at 37°C or 39°C for 24 h. DNAse agar (Britania) was used following product instructions to analyze extracellular DNAse activity. Extracellular protease was determined using TSA plates supplemented with 5% skimmed milk. Rhamnolipid production was analyzed in sheep-blood agar plates. Activity of extracellular lipases was analyzed using TSA plates supplemented with sterile egg-yolk suspension (5%). For all tests, the degradation halo or clear zone was measured in each case and the colony diameter was measured for normalization using the ImageJ software (65).

### Antibiotic and H_2_O_2_ sensitivity assays

Antibiotic sensitivity was determined using the commercial panel Sensititre (ThermoFisher) Gram positive for *S. aureus* USA300 and Gram negative for *P. aeruginosa* PAO1. The test was carried out strictly following the manufacturer’s instructions, but the plates were incubated at 37°C or 39°C. The MIC was determined following the instructions. H_2_O_2_ sensitivity was evaluated in agar plates as previously (66). Briefly, cultures grown as described in Figure S1 were seeded in a lawn, and a sterile Whatman no. 1 filter disc (6 mm) impregnated with 5 µM of 30% (v/v) H_2_O_2_ (Merck) was placed on the seeded plate. Plates were incubated only at 37°C since the H_2_O_2_ effect is almost instant. Inhibition zones were measured using the ImageJ software.

### Biofilm and motility assays

SA monocultures were seeded on Red Congo plates and then incubated overnight at 37°C or 39°C to detect exopolysaccharide production. Black-colored colonies were analyzed. Biofilm formation was analyzed in 96-multiwell polystyrene plates. Briefly, SA monocultures were inoculated at an initial OD_600nm_ of 0.025 in TSB supplemented with 0.5% KNO_3_. Incubation was carried out at 37°C or 39°C for 24 h. Planktonic cells were collected and OD_595nm_ was measured. Biofilm was stained with crystal violet as described before (67) and the attached biomass was determined at 550 nm in a plate reader (DR - 200Bs). For motility assays, the PA culture was adjusted to an OD_600nm_ of 1 (for co-cultures assay, the proportion was 1:1 for each bacterium) and a 5-µl drop was placed in a motility plate containing 8 g/l nutritive broth, 0.5% agar and supplemented with 0.5% glucose(68). Plates were incubated overnight at 37°C or 39°C. The entire movement was measured using the ImageJ software.

### Invasion assays on A549 cells

For invasion assays, A549 cells (a human adenocarcinoma epithelial cell line) were cultured at 37°C under 5% CO_2_ in Dulbecco’s Modified Eagle Medium (DMEM, Gibco) supplemented with 1% penicillin-streptomycin (Pen-Strep, Gibco) and 10% fetal bovine serum (FBS, Gibco). A549 cells were seeded in 24-well microtiter plates at 2.5 x 10^5^ cells per well and then incubated for 48 h (37°C, 5% CO_2_) in DMEM supplemented with 10% FBS and 1% Pen-Strep. Cells were washed twice with Dulbecco’s Phosphate-Buffered Saline (DPBS – Gibco) to remove antibiotics and then 1 ml DMEM supplemented with 10% FBS. Cells were infected at a multiplicity of infection (MOI) of 30 for SA monocultures and of 20 for PA monocultures. For invasion experiments with co-cultures, after determining the CFU/ml to the OD_600_ ratio of co-cultures, the OD_600nm_ was adjusted to 1. Then, 100 µl of this adjusted suspension was added to each well, which resulted in a MOI of 14 for SA and of 21 for PA. Bacteria and host cells were incubated for 1.5 h and then washed twice with DPBS. To kill extracellular bacteria, infected cells were incubated for 1.5 h in DMEM supplemented with 2.5 μg ml^−1^ lysostaphin (Sigma) for SA or gentamicin 200 μg ml^−1^ (Sigma) for PA infection or both for co-infections. Cells were washed again with DPBS and lysed with 0.1% Triton X-100, 0.5% trypsin, and 0.3 mg x ml^−1^ DNase (Sigma) in DPBS. Serial dilutions of cell lysates were plated in TSA to quantify the internalized bacteria for monocultures. In the case of co-infected cells, the lysates were plated on TSA-NaCl to detect SA and in cetrimide agar to detect PA. Results are presented as a percentage of the internalized bacterial inoculum.

### Cytokine measurements

For cytokine measurements, A549 cells were seeded in 96-well microtiter plates at a concentration of 5−10^4^ cells per well for 24 h (37°C, 5% CO_2_) in DMEM (Gibco) supplemented with 10% FBS (Gibco) and 1% Pen-Strep (Gibco). After that, cells were washed with DPBS, and then 198µl of fresh DMEM without antibiotics was added. Cells were stimulated by the addition of 2µl of bacterial cultures (OD_600nm_ adjusted to 0.1). After 24 h of incubation (37°C, 5% CO_2_), plates were centrifuged, and the supernatants were used for cytokine determinations. Human cytokine secretion was measured using the Invitrogen enzyme-linked immunosorbent assay (ELISA) kits for IL-6 and IL-8 according to the manufactureŕs instructions.

### Bacterial inoculum for mouse lung infection experiments

SA and PA were cultured as previously described. Cells from monocultures and co-cultures were incubated at 37°C or 39°C and then harvested by centrifugation at 8000 rpm for 5 min, washed twice using sterile PBS and resuspended in PBS. The resuspended cells were then used to infect mice intranasally. For SA, mice were infected with 2−10^8^ CFU per individual, whereas for PA infection, mice were infected with 1−10^7^ CFU per individual. For co-cultures, the resuspension was adjusted to an OD_600_ of 0.1 (corresponding to a concentration of 1.4 10^7^ CFU/ml for SA and 2.1 10^7^ CFU ml^−1^ for PA) and each mouse was infected with 200 µL.

### Mouse survival assay

Female DDY mice (6-8 weeks; 10 per group) were infected with SA monocultures at 37°C and 39 °C, PA monocultures preincubated at 37 °C and 39°C, or SA and PA co-cultures at 37 °C and 39 °C. Mice infected with sterile PBS were used as negative control. Mouse survival was monitored daily for 10 days. All experiments were carried out following the ethics guidelines and approved by the Institutional Ethics Committee of the University of Surabaya, Indonesia.

### Bacterial burden assay

For the bacterial burden assay, six mice per group were infected using the same bacterial growth conditions as in the mouse survival assays. Mice were sacrificed 24 h after infection, and the lungs, kidney and liver were aseptically recovered. These organs were then homogenized in sterile PBS containing 0.1% Tween 20. The bacterial load was determined by diluting the homogenized organ accordingly and plating on Mannitol Salt Agar (Merck) to count SA and Cetrimide Agar (Merck) to count PA. The CFU/ml were determined after 24 h incubation at 37 °C.

### Bacterial count in the presence of human serum albumin (HSA)

Bacterial count was performed in TSB medium supplemented or not with HSA (3.5%) following the same scheme described above. Cultures were plated in TSA+NaCl and cetrimide agar plates for SA and PA determinations, respectively.

### Statistical analysis

One- or two-way ANOVA with multiple comparisons or Unparied t-Test test was performed depending on the experiment, using GraphPad software (https://www.graphpad.com/). PCA was performed through the Singular Value Decomposition method on the normalized read counts. For multifactorial test applied to the entire RNAseq data set, two-way ANOVAs were carried out for each expression feature (genes) with factors: culture temperature (37° and 39° C), type (mono and co-culture), and interaction. The p-values of the interaction terms were adjusted for multiple testing with the Benjamini-Hochberg method, the same used by Rockhopper, which will be subsequently called q-values for consistency with this tool. PCA and multiple ANOVAs were carried out in R programming language (69).

### Ethics statement

Mouse experiments using Female DDY mice (6-8 weeks) were carried out with the approval of the Institutional Ethics Committee, University of Surabaya, Indonesia, no. 108/KE/IX/2022, following all the ethics guidelines.

## Author contributions

ESV (under the PMT and FG supervision) and PMT performed the conceptualization. ESV, MMR, MBG, SAR, DEE and PMT performed most of the investigation. AB and THM (under the supervision of AL) conducted the mice studies. NB and JE (under the supervision of MSR) carried out the HSA protection experiments. SN performed a formal multivariant analysis. FS, MMR and DDP carried out data curation. PMT and ESV wrote the manuscript. FG, AL and MMR reviewed and edited the manuscript. MMR, PMT and ESV carried out the visualization. PMT performed mostly the project administration and funding acquisition. MSR, DDP, AL and FG contributed with resources.

## Acknowledgments

we are thankful to Dr. Nancy I. Lopez for the proofreading of the manuscript and to Dr. Libera Lo Presti for editing. PMT discloses support for the research of this work from the ANPIDyI PICT 2018 N°2017 and by the Alexander von Humboldt Foundation (Equipment subsidy and Return Fellowship). FG discloses support for the research of this work from Cluster of Excellence EXC 2124 - Controlling Microbes to Fight Infections - 390838134. MSR discloses support for the research of this work from the National Institutes of Health (SC3GM125556). AL discloses support for the research of this work from Penelitian Keilmuan ITS 2022 (1022/PKS/ITS/2022). PMT, MMR and DDP are career investigators from CONICET AL holds an Alexander von Humboldt. MBG, SAR and FS have a doctoral fellowship from CONICET.

