## Supplemental Figures for "Fever-like temperature impacts on *Staphylococcus aureus* and *Pseudomonas aeruginosa* interaction, physiology, and virulence both *in vitro* and *in vivo*"

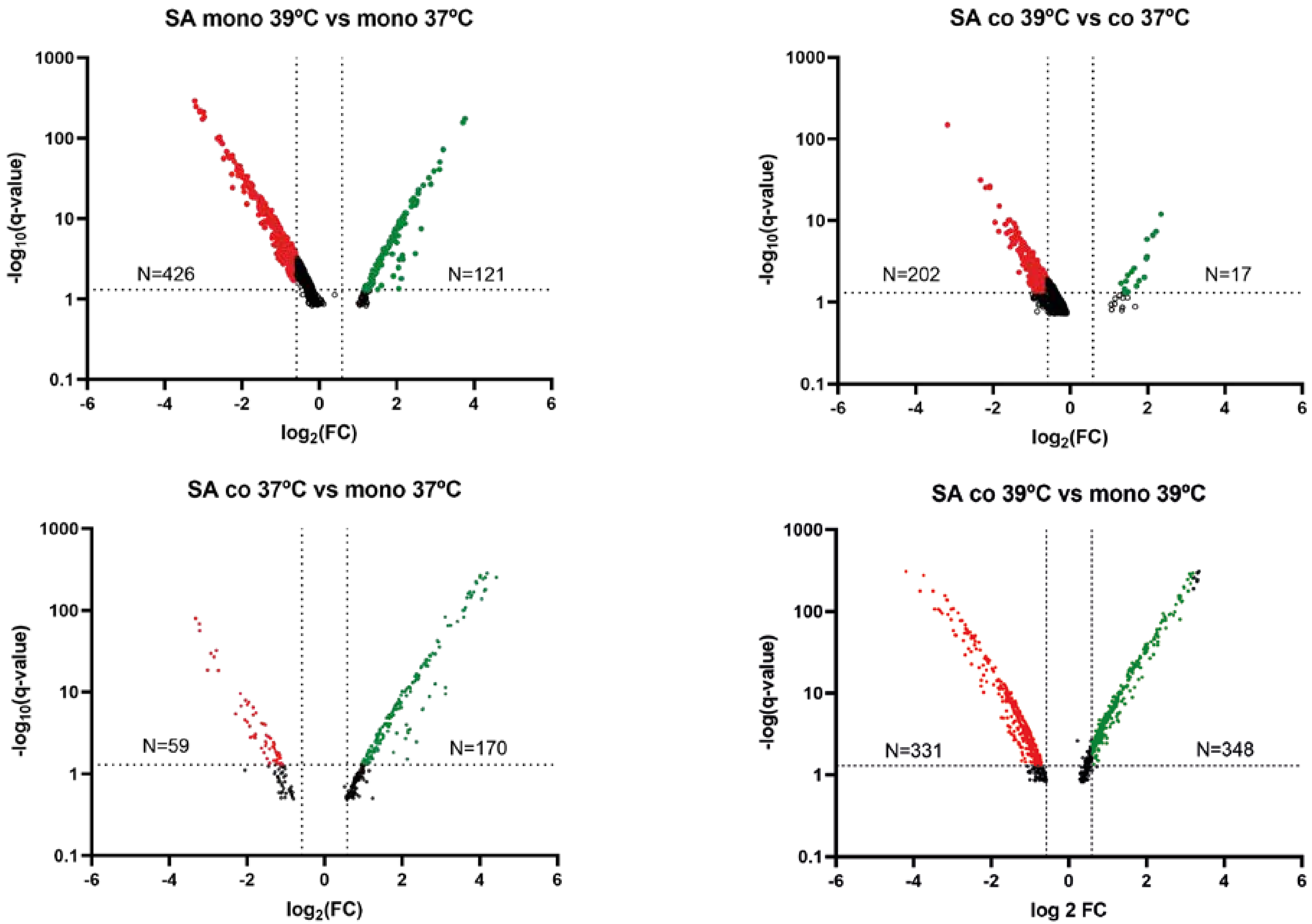


Fig S1: Volcano plots depicting –log10 (Q-value) versus log2 (Fold Change -FC-) for SA genes under different culture conditions (Fig.1). Differentially expressed genes (DEGs) are represented by colored dots. Green dots represent upregulated genes (P-value and Q-value <0.05, Fold change >1.5) and red dots represent downregulated genes (P-value and Q-value <0.05, Fold change < -1.5). For the construction of the Volcano plots, genes were initially filtered based on their P-values. All resulting genes were plotted

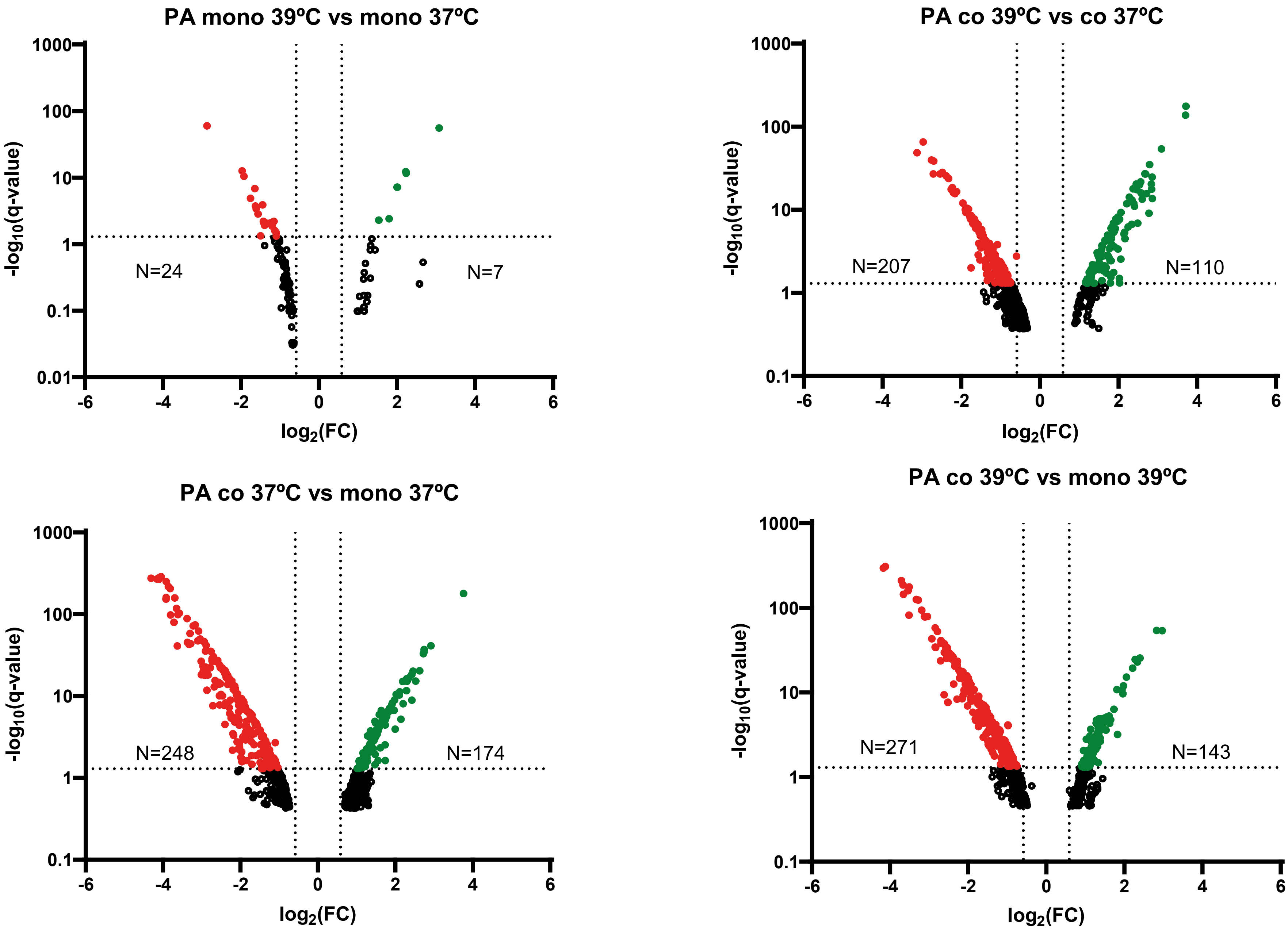


Fig S2: Volcano plots depicting –log10 (Q-value) versus log2 (Fold Change -FC-) for PA genes under different culture conditions (Fig.1). Differentially expressed genes (DEGs) are represented by colored dots. Green dots represent upregulated genes (P-value and Q-value <0.05, Fold change >1.5) and red dots represent downregulated genes (P-value and Q-value <0.05, Fold change < -1.5). For the construction of the Volcano plots, genes were initially filtered based on their P-values. All resulting genes were plotted

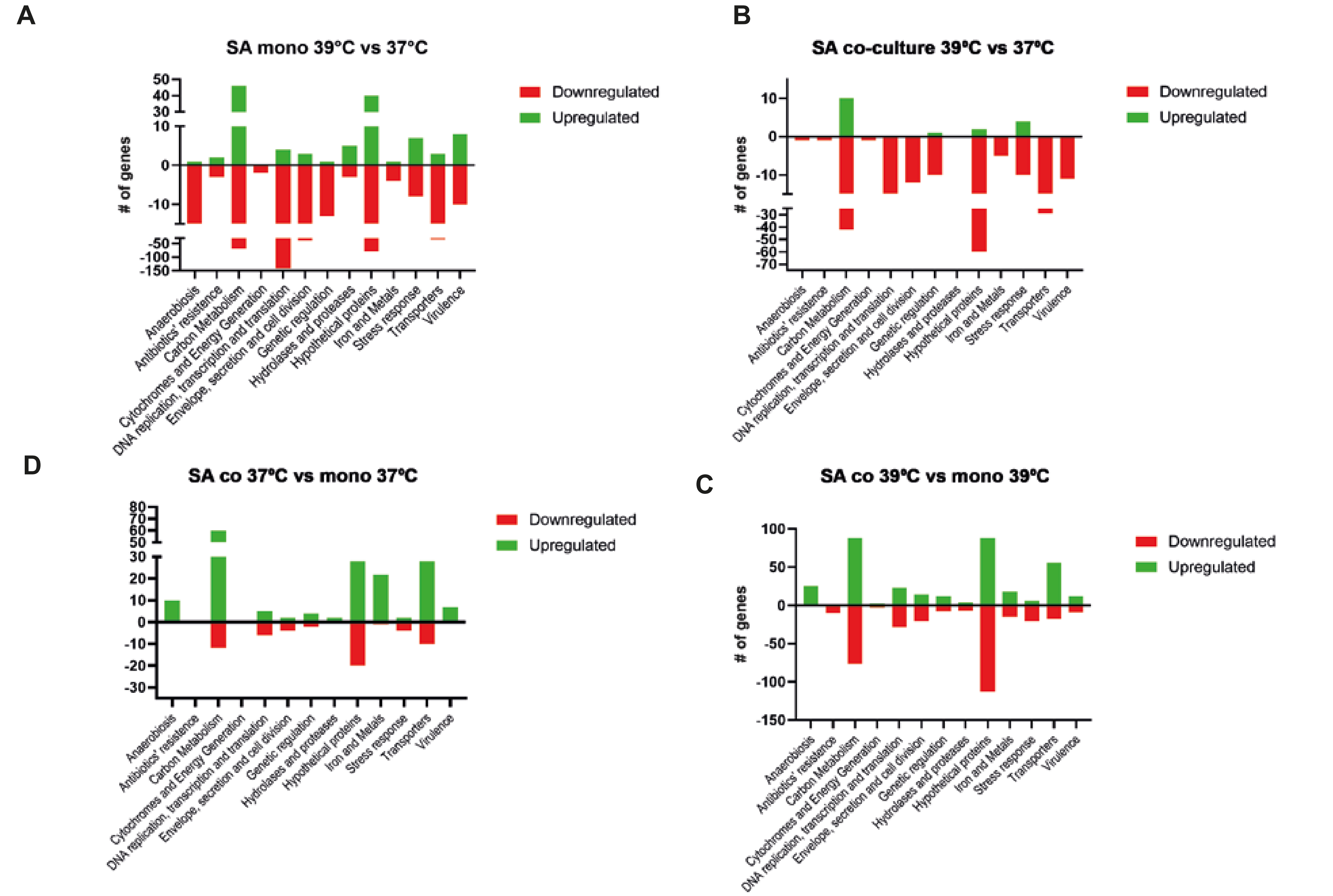


Fig S3: Functional classification for SA incubated under different culture conditions as described in Fig.1. Upregulated genes are represented in green while downregulated genes are shown in red.

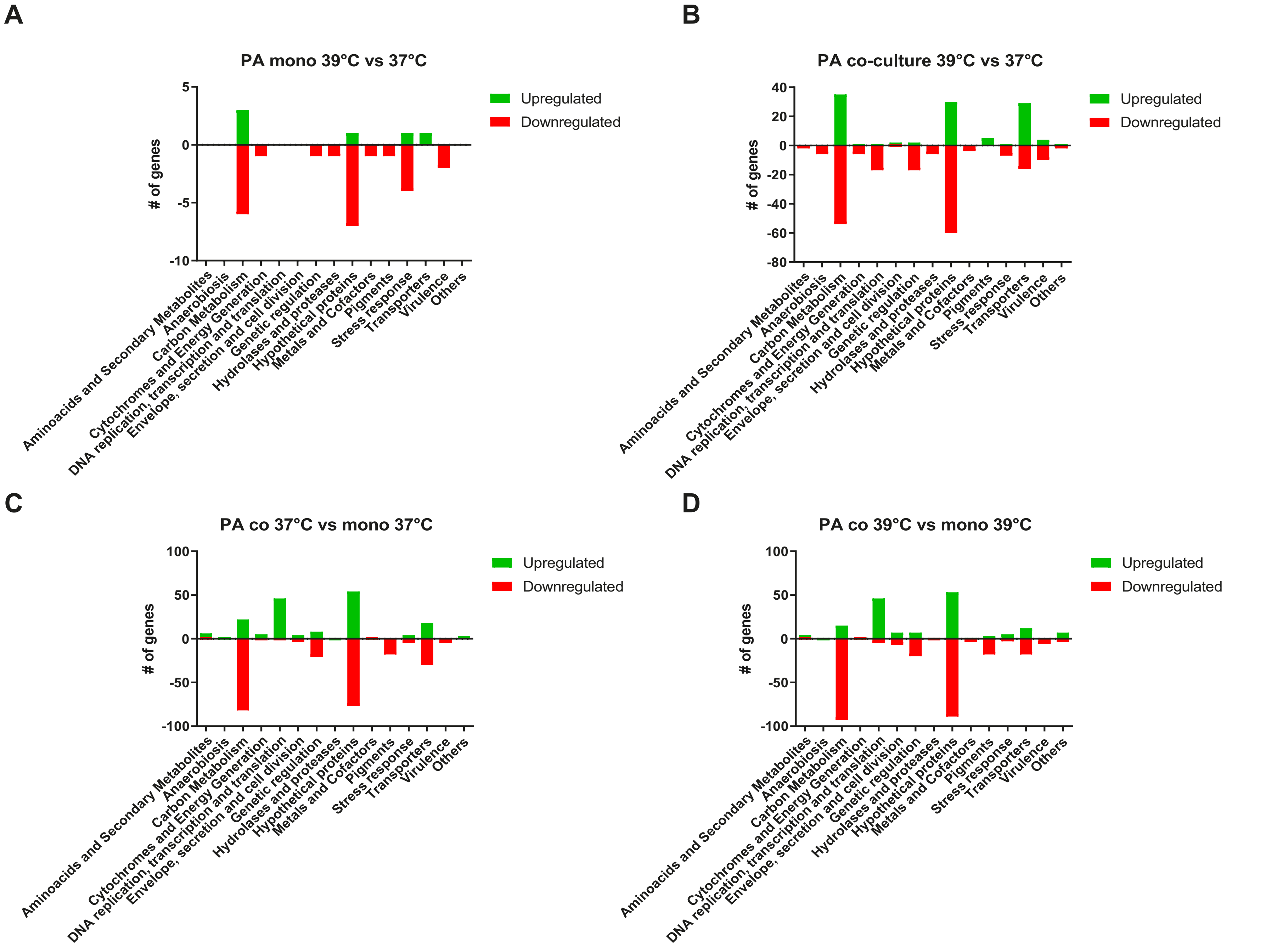


Fig S4 Functional classification for PA incubated under different culture conditions as described in Fig.1. Upregulated genes are represented in green while downregulated genes are shown in red.

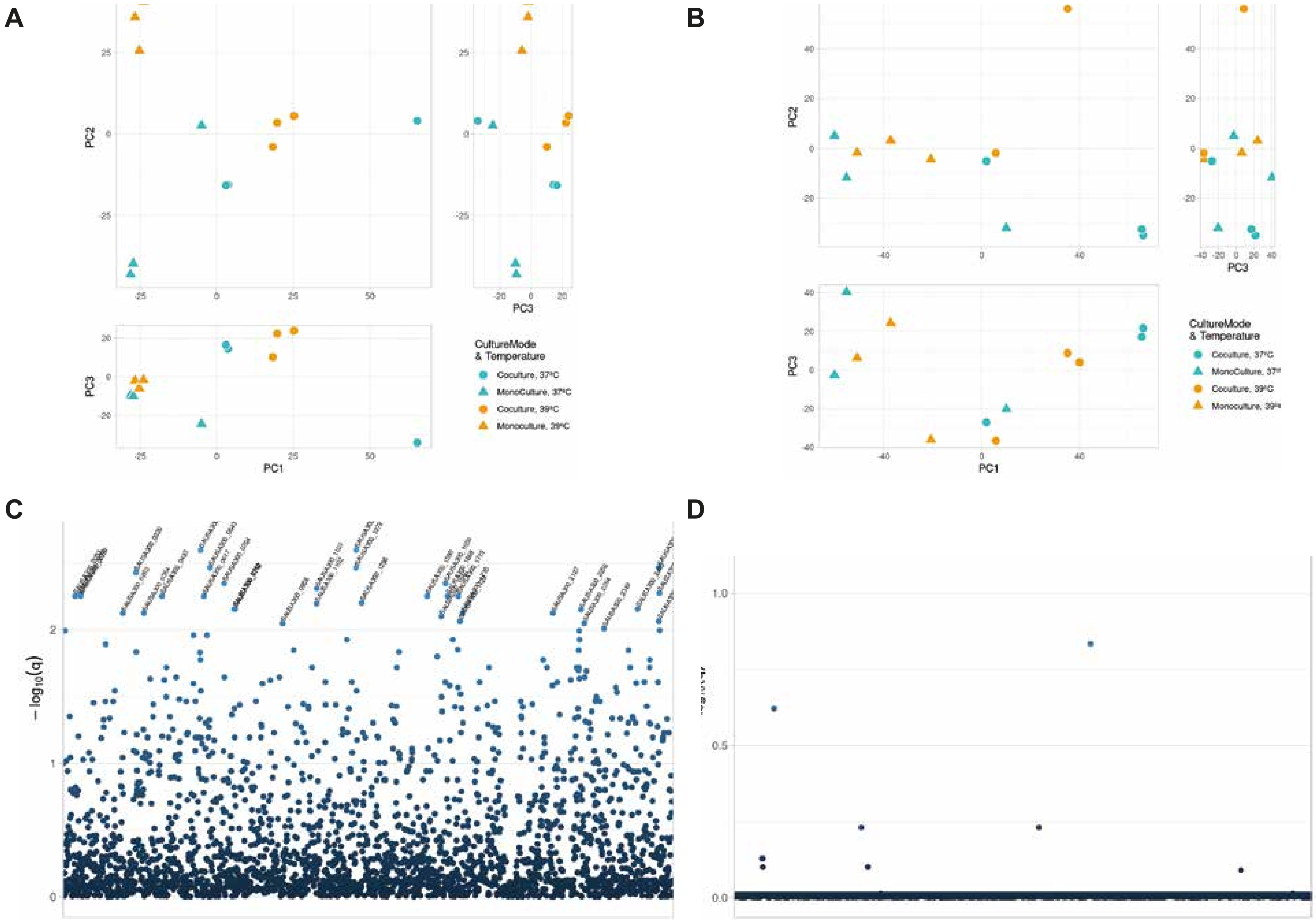


Fig S5: A. Principal Component Analysis for SA expression profile. B. Principal Component Analysis for PA expression profile. C. Interaction ANOVA for SA expressed genes D. Interaction ANOVA for PA expressed genes. Analysis was performed using R language.

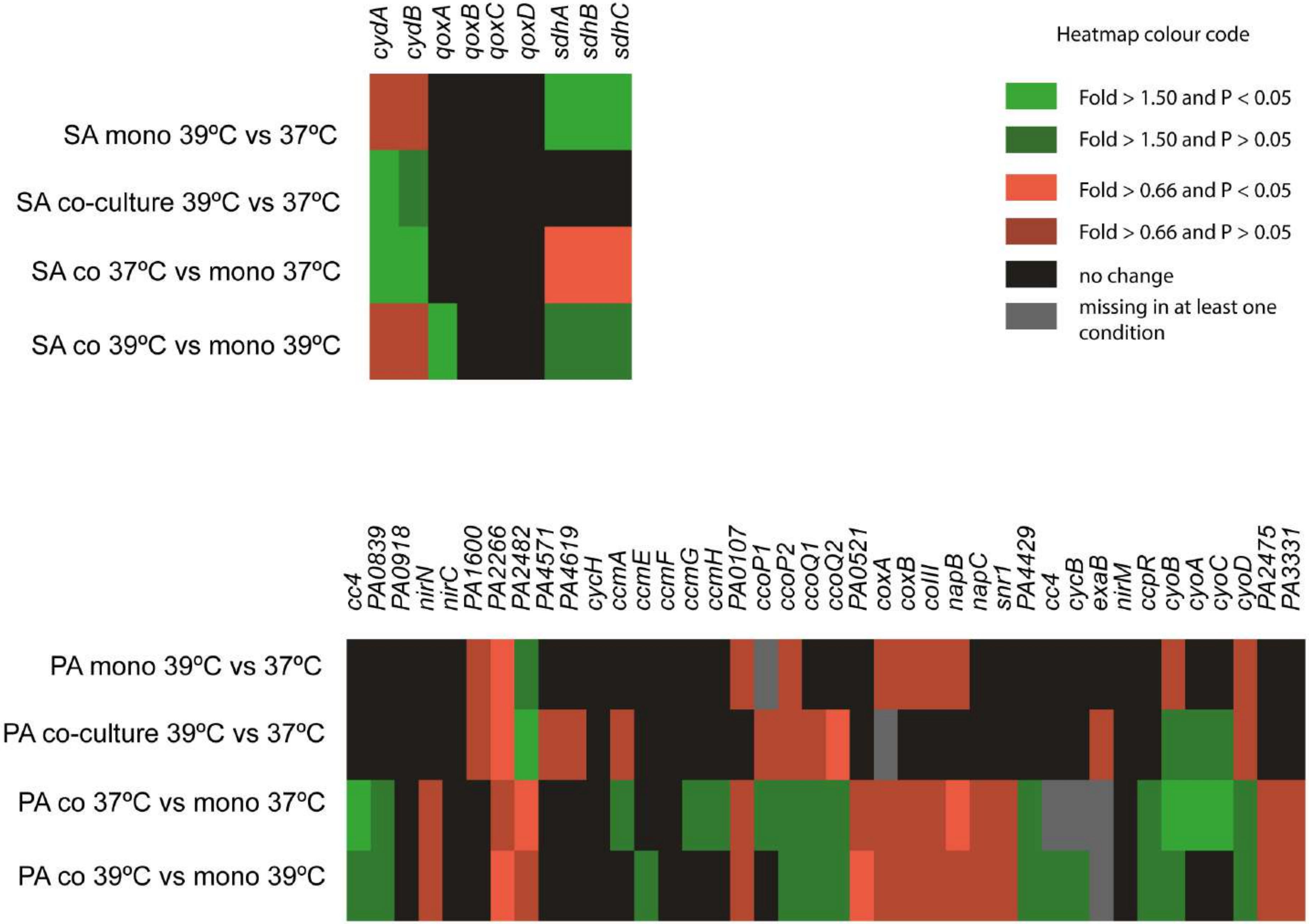


Fig S6: A. Cytochrome expression for SA. B. Cytochrome expression for PA.

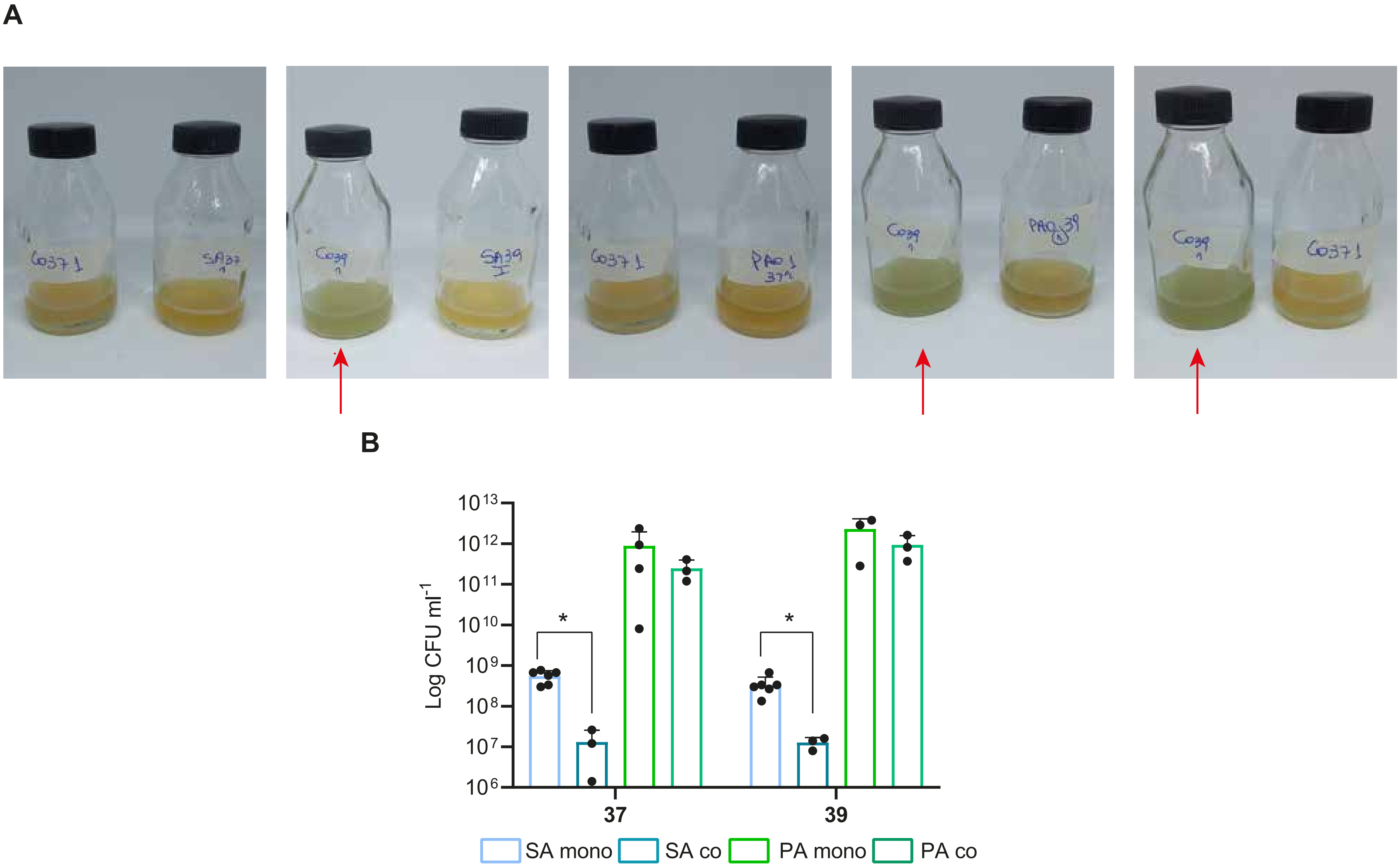


Fig. S7: A. SA, PA or cocultures incubated for 24 h at 37ºC or 39ºC. Pigment production in cocultures at 39ºC is marked with an arrow. B. SA CFU/ml and PA CFU/ml counting in monocultures and cocultures incubated at 37ºC or 39ºC for 24 h. * denotes significant differences.Even though data is displayed in the same graph, the following comparisons were performed independently using 1-way Anova: SA mono 37ºC vs SA co 37ºC P = 0.0008, SA mono 39ºC vs SA co 39ºC P = 0.0311
