## Supplemental Tables S3-S8 for "Fever-like temperature impacts on *Staphylococcus aureus* and *Pseudomonas aeruginosa* interaction, physiology, and virulence both *in vitro* and *in vivo*"

**Table S3:** Gene Ontology Analysis for enriched pathways in *S. aureus* expression profile

| **Enriched GEO pathways** | | | |
|---|---|---|---|
| **Monoculture 39 vs 37** | **Cocultures 39 vs 37** | **Coculture 37 vs monoculture 37** | **Coculture 39 vs monoculture 39** |
| Urea cycle | Superpathway of tetrahydrofolate biosynthesis and salvage | L-leucine biosynthesis | L-glutamate biosynthesis I |
| L-arginine degradation VI (arginase 2 pathway) | Tetrahydrofolate salvage from 5,10-methenyltetrahydrofolate | L-isoleucine biosynthesis I (from threonine) | L-isoleucine biosynthesis I (from threonine) |
| L-histidine degradation I | 5-aminoimidazole ribonucleotide biosynthesis | Superpathway of branched chain amino acid biosynthesis | Superpathway of branched chain amino acid biosynthesis |
| L-arginine biosynthesis II (acetyl cycle) | Inosine-5'-phosphate biosynthesis I | L-glutamate biosynthesis I/III | |
| L-arginine degradation I (arginase pathway) | UMP biosynthesis I superpathway of pyrimidine deoxyribonucleotides de novo biosynthesis | L-glutamine degradation I/ II | L-glutamine degradation I |
| Glycine cleavage | Glycine cleavage | L-serine degradation | L-serine degradation |
| Formate assimilation into 5,10-methylenetetrahydrofolate | Superpathway of pyrimidine ribonucleotides *de novo* biosynthesis | L-asparagine biosynthesis III (tRNA-dependent) | Pyridoxal 5'-phosphate biosynthesis II |
| Triacylglycerol degradation | Triacylglycerol degradation I | L-alanine degradation IV | L-threonine degradation |
| Flavin biosynthesis I | Inosine 5'-phosphate degradation | L-homoserine biosynthesis | |
| 2-oxoglutarate decarboxylation to succinyl-CoA | Guanosine nucleotides degradation III | Pyruvate fermentation | Pyruvate fermentation I/V |
| 5-aminoimidazole ribonucleotide biosynthesis I | di-trans poly-cis-undecaprenyl phosphate biosynthesis | ethanol degradation I | Ethanol degradation I/II |
| Gluconeogenesis I | Mevalonate degradation | Heme degradation VI | Heme degradation VI |
| Folate polyglutamylation | Molybdenum cofactor biosynthesis | Ammonia assimilation cycle III | Glycerol and glycerophosphodiester degradation |
| TCA cycle I | Biotin-carboxyl carrier protein assembly | Glutaminyl-tRNA-biosynthesis via transamidation | Heterolactic fermentation |
| Glycerol degradation V | UTP and CTP dephosphorylation folate transformations III | | Pyruvate fermentation to lactate |
| Mixed acid fermentation | Tetrapyrrole biosynthesis I (from glutamate) | | Mixed acid fermentation |
| Staphyloxanthin biosynthesis | Lipoteichoic acid biosynthesis | staphyloferrin B biosynthesis | Siroheme biosynthesis |

**Table S4:** Gene Ontology Analysis for enriched pathways in *P. aeruginosa* PAO1 expression profile

| Enriched GEO pathways | | | |
|---|---|---|---|
| **Monoculture 39 vs 37** | **Cocultures 39 vs 37** | **Coculture 37 vs monoculture 37** | **Coculture 39 vs monoculture 39** |
| Adenosyl-L-methionine salvage II | Alanine biosynthesis II | Alanine biosynthesis III | Isopropylamine degradation |
| Mixed acid fermentation | 2-oxoisovalerate decarboxylation to isobutanoyl-CoA | ATP biosynthesis | ATP biosynthesis |
| Glycerol degradation I | Biotin-carboxyl carrier protein assembly | Guanosine ribonucleotides *de novo* biosynthesis | tRNA charging |
| Glycerophosphodiester degradation | Branched-chain fatty acid biosynthesis | Heme biosynthesis II (oxygen-independent) | L-serine biosynthesis I |
| L-methionine biosynthesis | Superpathway of acetate utilization and formation | PRPP biosynthesis | L-tryptophan biosynthesis |
| Reactive oxygen species degradation | Glycine betaine degradation I | Biosynthesis from glutamate | L-phenylalanine biosynthesis |
| Glyoxylate bypass | Folate polyglutamylation | L-arginine degradation IX | Chorismate biosynthesis I |
| D-gluconate degradation | Superpathway of L-serine and glycine biosynthesis I | Arginine:pyruvate transaminase pathway | 3-dehydroquinate biosynthesis I |
| Fatty acid biosynthesis initiation (type II) | Pyocyanin biosynthesis | Selenate reduction | Selenate reduction |
| Thiosulfate oxidation III | 2-methylcitrate cycle I | | Sulfate activation for sulfonation |
| | Octane oxidation | | Assimilatory sulfate reduction IV |
| | Stearate biosynthesis II | | Lipopolysaccharide biosynthesis |
| | Glycine biosynthesis I | | Gluconeogenesis I |
| | Palmitate biosynthesis | | Polyhydroxydecanoate biosynthesis |
| | L-tyrosine degradation I | | tRNA processing |
| | L-valine degradation I | | Spermidine biosynthesis |
| | | | Coenzyme A biosynthesis I |
| | | | Lipid A biosynthesis I |

Table 1: Metabolic features of *S. aureus*

| Metabolic Pathway/Branch | Up-regulated genes | Down-regulated genes | Genes with increased expression (non-significative differences) | Genes with decreased expression (non-significative differences) |
|---|---|---|---|---|
| **Mono 39 vs 37** | | | | |
| **Arginine metabolism** | *arg-G, argH, rocD, gudB* | | *argJ, rocF, putA* | *argC, argF* |
| **TCA** | *mqo, gltA, icd, sucA, sucB* | | *sucC, sdhA sdhB* | |
| **Glycolysis** | | *plkA, fbaA* | | |
| **Peripheral feeding pathways** | *gudB, hutG* | | *hutH, hutI, hutU, rocA putA* | |
| **Fermentative metabolism** | *fdh* | *ldhA, pflAB, ackA pta, budA, budB* | | *gltA, gltB* |
| **Nitrate reduction** | | *narK, narHIJ, nirBD, nirR, nreABC* | | |
| **Staphyloxantin biosynthesis** | *crtN, crtM, crtP, crtQ crtO* | | | |
| **Co vs mono 37** | | | | |
| **L-isoleucine, L-valine and L-leucine** | *ilvA, ilvB , ilvC , ilvD, ilvN, leuA leuB* | | | |
| **Fermentative metabolism** | *ldhA, pflA, pflB* | | | |
| **Staphyloferrin B biosynthesis** | *sfnaA, sfnaB, sfnaC ,sfnaD* | | | |
| **Co vs mono 39** | | | | |
| **Glycolysis** | *glk, pfka, tpiA, gap, pgk, pgm, pyk* | | *pgi, fbaA* | |
| **TCA** | | | | |
| **Cytochrome** | | *sdhC, sdhA, sdhB* | | |
| **Staphyloferrin B biosynthesis** | *sfnaA, sfnaB, sfnaC, sfnaD* | | | |
| **Co 37 vs 39** | | | | |
| **Fermentative metabolism** | | *ldh, budB,* | | |

Table 2: Relevant metabolic features in *P. aeruginosa*

| Metabolic Pathway/Branch or cellular function | Up-regulated genes | Down-regulated genes | Genes with increased expression (non-significative differences) | Genes with decreased expression (non-significative differences) |
|---|---|---|---|---|
| **Mono 39 vs 37** | | | | |
| **Periplasmic glucose oxidation** | | *kgut, kguk, kgD, kgnt, eda* | | |
| **Co vs mono 37** | | | | |
| **Peripheral fructose catabolic pathway** | | *fruA, fruK fruI* | | |
| **Assimilatory sulfonate reduction** | *cysD, cysN, cysH,* alkane monoxygenase | | | |
| **L-lactate oxidation** | *lldD, lldP* | | | |
| **Co vs mono 39** | | | | |
| **Periplasmic glucose oxidation pathway** | | *gdc, kgut, kguK, kguD, glk* | | |
| **Glyoxylate shunt** | *icl, sucC, sucD* | | | |
| **Lactate oxidation** | *lldD, lldP* | | | |
| **Co 39 vs 37** | | | | |
| **Ethanol oxidation** | *pqqC, pqqD, exaC, erbR* | | *pqqA, pqqB, pqqE, erbS* | |
| **Anaerobic metabolism** | | *arcD, nrdD, narK1, narK2, dnr, aer2, hcnB* | | |

**Table 1:** genes related to virulence that present in its expression interaction between temperature and culture condition in SA using Multifactorial Anova with Benjamini-Hochberg method.

| **Locus tag** | **Gen** | **Product** | **Q interaction** |
|---|---|---|---|
| SAUSA300_1989 | *agrB* | accessory gene regulator protein B | 0.0443833 |
| SAUSA300_1991 | *agrC* | accessory gene regulator protein C | 0.0444769 |
| SAUSA300_1992 | *agrA* | accessory gene regulator protein A | 0.0458893 |
| SAUSA300_1067 | *psmβ1* | anti protein | 0.0374364 |
| SAUSA300_1068 | *psmβ2* | anti protein | 0.0389214 |
| SAUSA300_1988 | *hld* | delta-hemolysin | 0.0341659 |

**Table S5:** Antibiotic sensitivity for *S. aureus* in Sensitrine ARGPF plates incubated at different temperatures.

| **Antibiotic** | **MIC (µg/ml)** | |
|---|---|---|
| | 37$^{o}$C | 39$^{o}$C |
| Vancomycin | 1 | 1 |
| Chloramphenicol | 8 | 8 |
| Penicillin | >8 | >8 |
| Rifampicin | ≤0.5 | ≤0.5 |
| Ampicillin | >8 | >8 |
| Tigecycline | 0.25 | 0.25 |
| **Moxifloxacin** | **2** | **4** |
| Erythromycin | >4 | >4 |
| oxacilin+2%NaCl | >4 | >4 |
| Levofloxacin | >4 | >4 |
| Nitrofurantoin | ≤32 | ≤32 |
| **Daptomycin** | **≤0.5** | **1** |
| Linezolid | 2 | 2 |
| Ciprofloxacin | >4 | >4 |
| Tetracycline | ≤2 | ≤2 |
| Gentamicin | ≤2 | ≤2 |
| Minocycline | ≤4 | ≤4 |
| Clindamycin | ≤0.5 | ≤0.5 |
| Streptomycin 1000 | NEG | NEG |
| Gentamicin 500 | NEG | NEG |

**Table S6:** Antibiotic sensitivity for *P. aeruginosa* PAO1 in Sensitrine ARGNP plates incubated at different temperatures

| **Antibiotic** | **MIC (µg/ml)** | |
|---|---|---|
| | 37$^{o}$C | 39$^{o}$C |
| Ceftazimide | ≤2 | ≤2 |
| Ceftazimide/clavunalic acid | 0,5/4 | 0.5/4 |
| Ceftaximide/clavunalic acid | >2/4 | >2/4 |
| **Cefotaxime** | **32** | **8** |
| Ampicilin/sulbactam (2:1 ratio) | >16/8 | >16/8 |
| Ampicillin | 8< | 8< |
| Piperacilin/tazobactam constant 4 | ≤8/4 | ≤8/4 |
| Cepefime | ≤2 | ≤2 |
| Meropenem | ≤1 | ≤1 |
| Cephalothin | >32 | >32 |
| Iminipem | 2 | 2 |
| Amikacin | ≤8 | ≤8 |
| Levofloxacin | 4 | 4 |
| Gentamycin | ≤4 | ≤4 |
| Ciprofloxacin | >2 | >2 |
| **Minocycline** | **8** | **>8** |
| Tigecycline | >2 | >2 |
| Cefuroxime | >16 | >16 |
| Ertapenem | >2 | >2 |
| Colistin | ≤1 | ≤1 |
| Cefoxitin | >16 | >16 |
| Doripenem | ≤4 | ≤4 |
| Rifampin | >8 | >8 |
| Nitrofurantoin | >64 | >64 |
| Fosfomycin+glucose 6 phosphare | >fos+ 64 | >fos+ 64 |
| Trimethoprim/sulfamethoxazole | >2/38 | >2/38 |
| Chloramphenicol | >16 | >16 |
| Amoxicilin/clavulanic acid 2:1 ratio | >16/8 | >16/8 |
| Aztreonam | ≤8 | ≤8 |
| Nalidixic acid | >16 | >16 |
